# *Ecdysone Receptor* autonomously controls germ cell differentiation in the *Drosophila* ovary

**DOI:** 10.64898/2025.12.19.695538

**Authors:** Lauren E. Jung, Alexandria I. Warren, Changhong Yin, Weihua Huang, Allison C. Simmons, Samantha I. McDonald, Lindsay A. Swain, Victoria E. Garrido, Daniel N. Phipps, BiClaireline Cesar, Danielle S. Finger, Zhipeng Sun, Todd G. Nystul, Elizabeth T. Ables

**Author notes:** These authors contributed equally to this work (Lauren E. Jung, Alexandria I. Warren). Correspondance: **Corresponding author:** Elizabeth T. Ables, East Carolina University Department of Biology, 1001 E. 10^th^ St., Mailstop 552, 553 Science & Technology Building Greenville, NC 27858.

## Abstract

In the *Drosophila* ovary, the steroid hormone ecdysone controls germline stem cell (GSC) maintenance and germ cell differentiation. Prior studies demonstrated that ecdysone regulates germ cell differentiation non-autonomously via the nuclear receptor Ecdysone Receptor (EcR) in ovarian somatic cells. Although EcR is also expressed in GSCs and their differentiating daughters, potential direct roles for EcR in GSCs independent of the soma have not been examined. Here, we demonstrate that EcR functions autonomously in GSCs and cystoblasts to control germline differentiation. While depletion of *EcR* from GSCs mildly reduces GSC self-renewal, over-expression of *EcR* specifically in GSCs and cystoblasts impedes germ cell differentiation, phenotypically resembling *bag of marbles* loss-of-function and Bone Morphogenetic Protein signaling constitutive activation. We propose that while low levels of EcR are essential to maintain GSC self-renewal and permit initial differentiation, higher levels of EcR accumulate in differentiated germ cells to promote transcription of maternal genes, providing temporal control over germline differentiation. These data support the model that stem cells harbor unique mechanisms to integrate signals from multiple cell sources that safeguard their self-renewal in response to local and physiological cues.

**SUMMARY STATEMENT:** The nuclear receptor EcR modulates stem cell maintenance and differentiation in ovarian germ cells.

## INTRODUCTION

Physiological signals provide spatiotemporal cues linking tissue-resident stem cell activity with tissue demand. For example, stem cells in the skin, muscle, mammary gland, and hematopoietic system respond to hormone fluctuations associated with puberty and pregnancy (Chhabra and Booth, 2021; de Morree and Rando, 2023; Hsu and Fuchs, 2022; Nakada et al., 2014). As the primary receptors for steroid hormones and other lipophilic signals, nuclear receptors serve as key molecular nodes that coordinate stem cell activity in response to physiological changes. Crosstalk between nuclear receptors influences cell plasticity and heterogeneity; moreover, dysregulation of nuclear receptor signaling contributes to many disease states (BharathwajChetty et al., 2024; Simandi et al., 2013; Weikum et al., 2018). Yet, despite the highly conserved nature of nuclear receptor structure and function, the molecular mechanisms by which nuclear receptors control stem cell homeostasis are not well-understood.

*Drosophila* ovarian germline stem cells (GSCs) are an excellent model to study the response of tissue-resident stem cells to nuclear receptors (Hinnant et al., 2020; Spradling et al., 2022). GSCs reside in the germarium (a niche at the anterior-most tip of adult ovaries) and provide the cellular progenitors necessary to create oocytes (Figure 1A). GSCs divide asymmetrically to produce a cystoblast, which undergoes four rounds of mitotic division with incomplete cytokinesis to produce an interconnected 16-cell cyst. Within the cyst, one cell differentiates as the oocyte while the remaining 15 become nurse cells, which produce maternal mRNA and proteins that are transported into the oocyte and necessary for embryonic development post-fertilization. Somatic cells surround GSCs and developing cysts and are integral to proper oocyte development and function.

**Figure 1.**
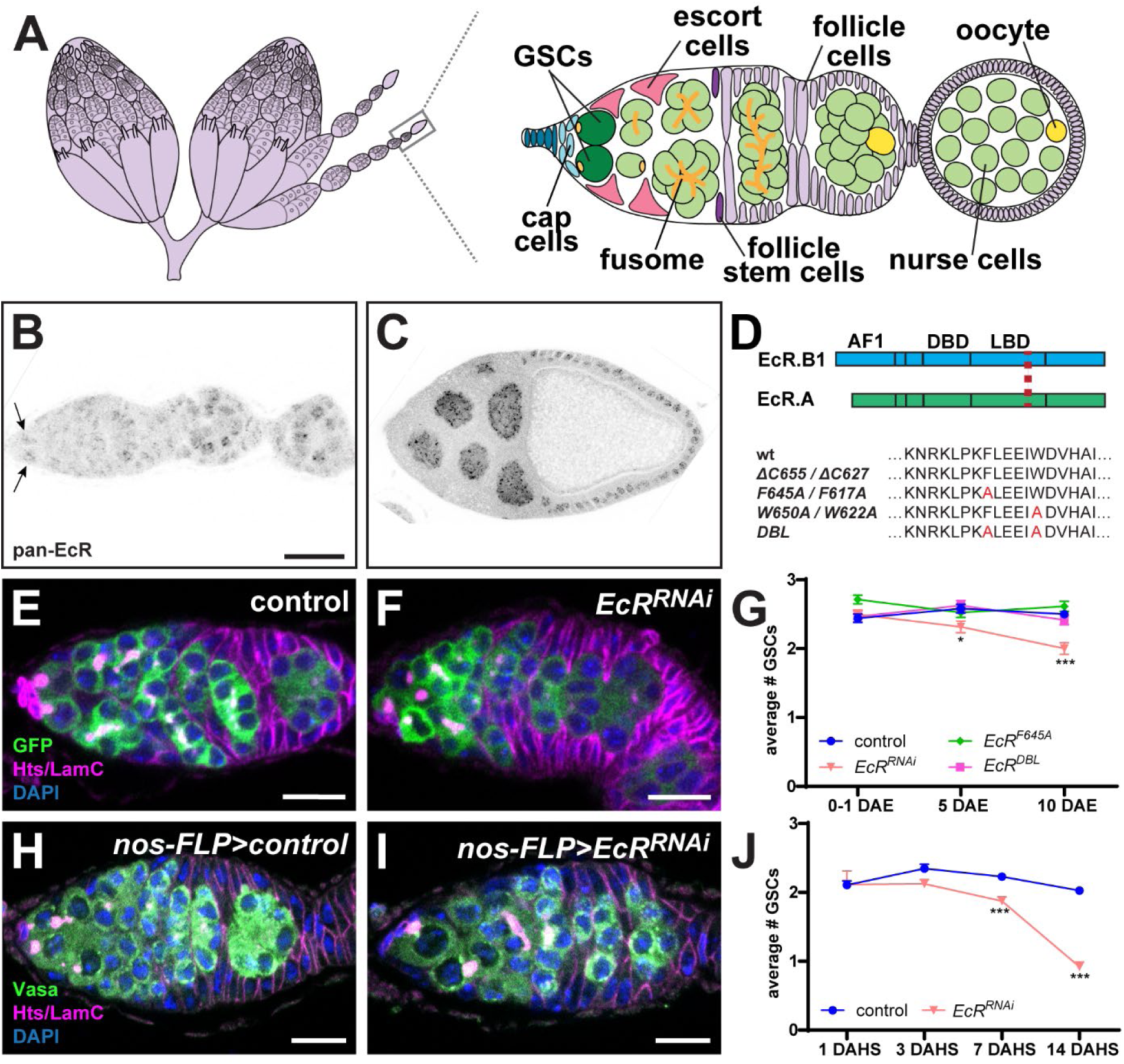
*EcR* is expressed in GSCs and necessary for their maintenance. **(A)** Schematic of a *Drosophila* ovary. The germarium is the anterior-most region of each ovariole (boxed area enlarged at right). GSCs (green) divide to create cystoblasts and cysts (pale green), featuring the fusome (orange), a germ cell-specific organelle. The oocyte (yellow) is selected from within the 16-cell cyst. Germ cells are surrounded by somatic cells: cap cells (light blue), escort cells (pink), follicle stem cells (purple) and follicle cells (pale purple.) **(B-C)** Immunostaining for EcR in wild-type ovariole. Arrows indicate GSCs. (C) is a stage 10 egg chamber from the same ovariole as (B), imaged at the same fluorescence intensity to compare expression levels. **(D)** Schematic comparison of EcR-B1 and EcR-A proteins, indicating the DNA binding domain (DBD) and ligand binding domain (LBD). Red text indicates amino acid substitutions in the LBD, listed for wildtype and dominant-negative transgenes below. **(E-G)** Representative images (E-F) and quantification (G) of GSCs per germarium in *nos-Gal4::VP16>UASp-GFP::tubulin* control (E) and *nos-Gal4::VP16>UASp-GFP, EcR^RNAi1^* germaria immunostained for GFP (green), Hts (magenta; fusomes and follicle cells), LamC (magenta; cap cells), and DAPI (blue; nuclei) over time. Quantification performed at 0-1, 5, and 10 days after eclosion (DAE); *n* > 100 germaria per genotype per timepoint. **(H-J)** Representative images (H-I) and quantification (J) of GSCs per germarium in *hsFLP;nos-STOP-Gal4* control (abbreviated *nos-FLP*; H) or *nos-FLP>EcR^RNAi1^* immunostained for Vasa (green), Hts and LamC (magenta), and DAPI (blue). Quantification performed at 1, 3, 7, and 14 days after heat shock (DAHS); *n* > 72 germaria per genotype per timepoint. Scale bars = 20µm (B-C) or 10µm (E-F,H-I). Error bars represent s.e.m., \**p*<0.01, \*\*\**p*<0.0001, Student’s t-test.

GSC proliferation and maintenance depend on ecdysone, a steroid hormone well-known to regulate developmental timing in insects (Ables and Drummond-Barbosa, 2010; Ables et al., 2016; Ameku and Niwa, 2016; König et al., 2011; Morris and Spradling, 2012; Shi et al., 2021). Ecdysone is synthesized from dietary cholesterol through a multi-step biosynthesis cascade, and ecdysone titer acts as a temporal cue to initiate developmental progression, pupariation, and the final metamorphic transition to adulthood (Morrow and Mirth, 2024). Ecdysone controls cellular function by binding the Ecdysone Receptor (EcR), a complex of two nuclear receptors (encoded by *EcR/NR1H* and *usp/NR2B*) that regulate gene expression (Morrow and Mirth, 2024; Uyehara and McKay, 2019). Cell type-specific responses are influenced in part by differential expression and transcriptional activity of three EcR protein isoforms (EcR-A, EcR-B1, and EcR-B2) sharing conserved DNA and ligand binding domains (Schauer et al., 2011; Schubiger et al., 2003; Talbot et al., 1993). The ovary is the primary source of ecdysone in adult females, and its titer is dependent on female age, reproductive status, and diet (Harshman et al., 1999; Hodgetts et al., 1977; Riddiford, 1993).

Global genetic loss-of-function studies indicate that ecdysone signaling promotes GSC self-renewal and cyst formation (Ables and Drummond-Barbosa, 2010; Morris and Spradling, 2012). While these processes are indirectly controlled by EcR activity in somatic cells (Ameku and Niwa, 2016; König and Shcherbata, 2015; König et al., 2011; Morris and Spradling, 2012; Shi et al., 2021), evidence suggests that EcR also functions autonomously in the early germline. First, germ cell-specific inactivation of the EcR obligate co-receptor *usp* blocked GSC self-renewal and cyst formation (Ables and Drummond-Barbosa, 2010). Second, several ecdysone-responsive genes are expressed in germ cells, including other nuclear receptors (Ables et al., 2015; Beachum et al., 2021; Buszczak et al., 1999; McDonald et al., 2019; Zike et al., 2025). Finally, germ cell-specific inactivation of putative EcR transcriptional target genes blocked GSC self-renewal (Ables et al., 2016). However, a definitive link between EcR and its putative transcriptional targets has not been established. This is a critical knowledge gap, since many ecdysone-responsive genes may also function independently of EcR (Liu et al., 2025).

Prior studies also suggest that EcR activity in the early germline is coordinated with spatially-regulated Bone Morphogenetic Protein (BMP) signaling (Ables and Drummond-Barbosa, 2010; Drummond-Barbosa, 2019). BMP ligands are secreted from somatic cap cells and received via Transforming Growth Factor beta-class Ser/Thr kinase receptors, including the Type-I receptor encoded by *thickveins* (*tkv*). Upon signal reception, the transcription factor encoded by *Mothers against decapentaplegic* (*Mad*) is phosphorylated and translocated to the nucleus. Mad represses transcription of *bag of marbles* (*bam*), an essential differentiation factor. Division of the GSC perpendicular to the niche places the cystoblast farther from BMP ligands, permitting transcription of *bam*. Mad also promotes transcription of *Daughters against decapentaplegic* (*Dad*), which aids in degradation of active Tkv protein (Liu et al., 2023). Over-expression of constitutively active Tkv (*tkv^ACT^*) or knock-down of *bam* in germ cells results in accumulation of undifferentiated germ cells (Casanueva and Ferguson, 2004; McKearin and Spradling, 1990; Xie and Spradling, 1998).

Here, we present evidence that EcR autonomously controls two steps of germline differentiation. EcR is expressed at low levels in GSCs and cystoblasts, and depletion of *EcR* via germline-enhanced RNAi is sufficient to reduce GSC self-renewal over time. In contrast, over-expression of *EcR* in GSCs and cystoblasts delays germ cell differentiation, creating a mixed population of undifferentiated, proliferative germ cells that phenocopy loss of *bam* or constitutively active BMP signaling. We find evidence suggesting that EcR and BMP signaling converge on a common set of transcriptional targets, including *tkv* itself and the transcription factor *longitudinals lacking* (*lola*). Moreover, EcR-regulated genes are enriched in *bam* mutant germ cells, suggesting that EcR plays a secondary role in cystoblasts and/or differentiating cysts. We propose that EcR functions independently in GSCs and differentiating germ cells, controlled by the level of receptor and the level of available hormone, to promote stem cell maintenance and activation of the maternal transcriptional program as cells differentiate. This activity, combined with the non-autonomous action of EcR in adjacent cap and escort cells, collectively modulates GSC retention in the niche and germ cell differentiation in response to physiological cues.

## RESULTS

### EcR is expressed at low levels in the germline that are necessary for GSC self-renewal

Although prior studies described ubiquitous expression of EcR and Usp in the ovary, available reporters of EcR activity are not expressed in germ cells, and Usp protein is present at much lower levels in GSCs than in the surrounding soma or in cysts (Buszczak et al., 1999; König et al., 2011; Morris and Spradling, 2012). We thus verified that EcR is expressed in the germline (Figure 1B-C). Using an antibody against all EcR isoforms, we observed nuclear EcR localization in cells throughout the germarium, including GSCs (Figure 1B). EcR levels were lower in undifferentiated germ cells than in 16-cell cysts, and lower in the germline compared to somatic follicle cells until stage 10 of oogenesis (Figure 1C).

We then asked whether depletion of *EcR* or inactivation of EcR signaling in germ cells was sufficient to inhibit long-term retention of GSCs in the niche. Using a germline-specific driver (*nos-Gal4::VP16*), we depleted *EcR* using a germline-enhanced RNAi (*EcR^RNAi1^*, which targets all *EcR* isoforms; Figure S1A-C) and inactivated EcR signaling by over-expressing new germline-permissive dominant-negative forms of *EcR* resulting from point mutations in the ligand binding domain [*EcR-B1^W650A^*, *EcR-B1^F645A^*, or *EcR-B1^F645A+W650A^* (referred to as *EcR^DBL^*); (Figure 1D and S1D-E)]. Prior *in vitro* studies demonstrated that both *EcR-B1^W650A^*and *EcR-B1^F645A^* fail to stimulate transcription, but *EcR-B1^F645A^*binds ligand normally; both mutants function as competitive inhibitors of wild-type protein (Cherbas et al., 2003; Hu et al., 2003). We counted GSCs (identified by their anteriorly-localized fusome) in germaria collected from age-matched female flies from eclosion to 10 days post-eclosion (Figure 1E-G). Although over-expression of the dominant-negative *EcR* alleles did not impact GSC number, depletion of *EcR* via RNAi reduced the number of GSCs over time (Figure 1G). Since the effect on GSC number could reflect alterations in either GSC establishment or maintenance, we depleted *EcR* specifically at adulthood using inducible RNAi (Figure 1H-J). Conditional depletion of *EcR* in germ cells post-eclosion similarly reduced GSC number as flies aged, suggesting EcR regulates GSC self-renewal, but not their establishment (Figure 1H-J).

### Over-expression of *EcR* in GSCs and cystoblasts delays germ cell differentiation in a ligand-dependent manner

Since EcR is expressed at low levels in GSCs and cysts, we asked whether the levels of EcR must be kept low to permit germ cell differentiation. We co-expressed wild-type *EcR-B1* with a fluorescently-tagged *α-Tubulin* (*UASp-α-Tub::GFP*) to label germ cells using a germline-specific driver (Figure 2A-B). As expected, over-expression of *α-Tubulin::GFP* alone yielded fully developed ovaries containing two or three GSCs, mitotically dividing cysts, and developing egg chambers (Figure 2A,J). In contrast, over-expression of *EcR-B1* resulted in female sterility. Ovaries were small and lacked fully-developed egg chambers. Germaria were full of undifferentiated singlet and doublet germ cells, characterized by a single round or stretched fusome (a germ cell-specific organelle) (Figure 2B,J). We also built two additional transgenes in which wild-type EcR was terminally tagged with a fluorophore (Figure 2C-D). Like the untagged version, expression of *mCherry::EcR-B1* blocked germline differentiation; however, expression of *EcR-B1::mCherry* blocked differentiation but at lower penetrance, allowing some egg chambers to differentiate (Figure 2C-D,J). This suggests that the C-terminal fluorophore causes a conformational change in the protein, reducing function. mCherry::EcR-B1 protein localized to both germ cell nuclei and cytoplasm, while EcR-B1::mCherry localized exclusively to nuclei (Figure 2C-D).

**Figure 2.**
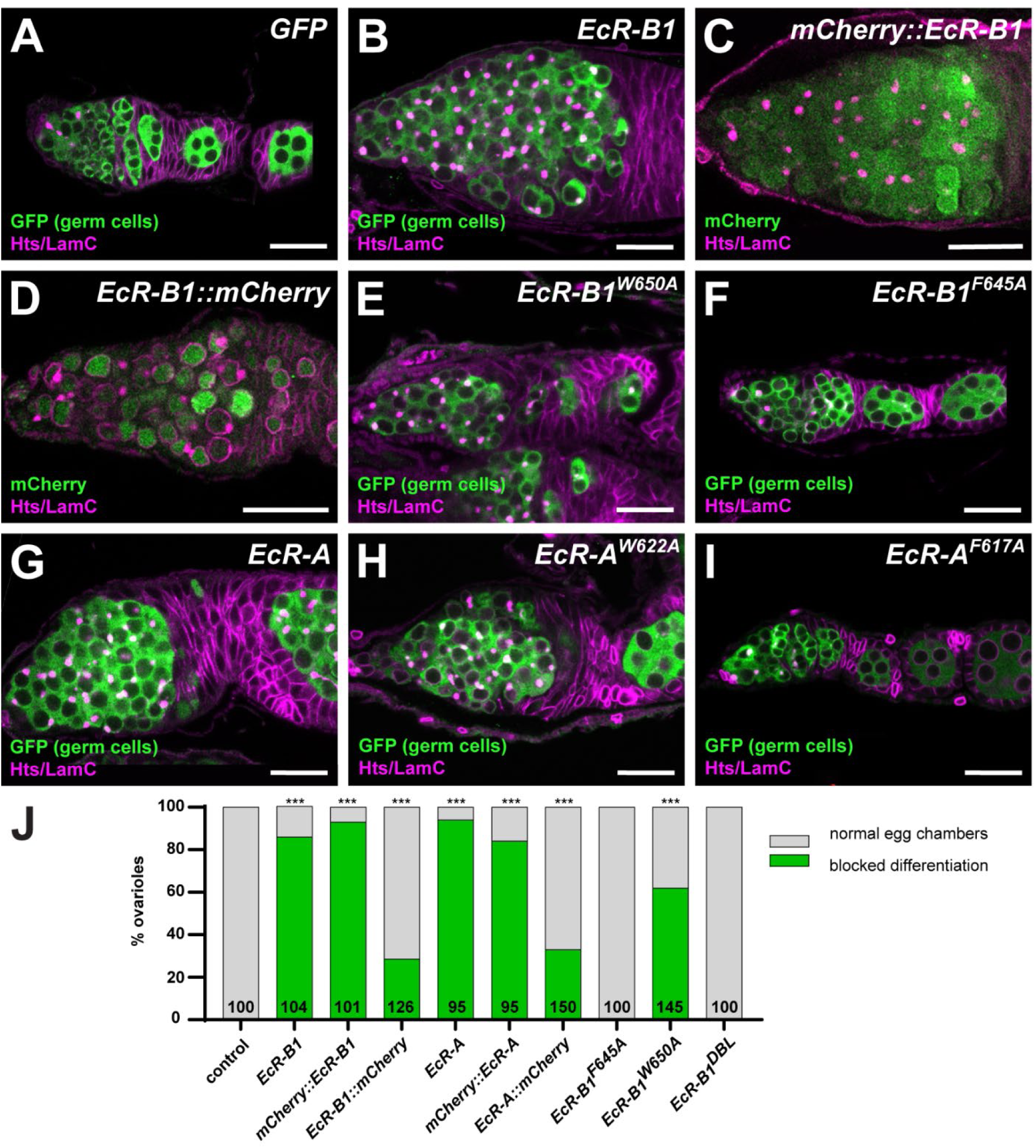
Over-expression of *EcR* in germ cells impairs their differentiation. **(A-J)** Representative images (A-I) and quantification (J) of *nos-Gal4::VP16* driving *UASp-GFP::tubulin* alone (A, control) or with *UASz-EcR-B1* (B), *UASz-mCherry::EcR-B1* (C), *UASz-EcR-B1::mCherry* (D), *UASz-EcR-B1^W650A^* (E), *UASz-EcR-B1^F645A^* (F), *UASz-EcR-A* (G), *UASz-EcR-A^W622A^* (H), or *UASz-EcR-A^F617A^* (I) germaria immunostained for GFP (green, germ cells, A-B,E-I) or mCherry (green, transgene expression, C-D), Hts (magenta; fusomes and follicle cells), and LamC (magenta; cap cells and stalk cells). Scale bars = 20 µm. Bars in (J) represent the percentage of ovarioles displaying a block to germ cell differentiation (green); *n* > 50 ovarioles from at least 15 flies per genotype, ∼5 days after eclosion. \*\*\**p*<0.0001, compared to control; Fisher’s exact test.

We then tested whether *EcR-B1* over-expression specifically in dividing cysts (exclusive of GSCs) could block germ cell differentiation. Using three independent transgenes that drive *Gal4* expression in dividing cysts (*3xbam-Gal4*), dividing cysts and 16-cell cysts (*aub-Gal4*), or post-mitotic cysts (*otu-Gal4::VP16*), we discovered that restricting over-expression of *EcR-B1* to the mitotic or post-mitotic cysts is not sufficient to block their differentiation (Figure S2A-F). These results suggest that increased titer of EcR protein in dividing cysts does not impair their division or differentiation, consistent with the endogenous increase in EcR levels as cysts divide.

Because EcR can impact transcription in the presence or absence of ligand (Uyehara et al., 2022), we asked whether EcR activity in early germ cells reflected an ecdysone-dependent function of the receptor. We first quantified germline differentiation in females that over-expressed the germline-permissive dominant-negative forms of *EcR* (Figure 1D). While expression of *EcR-B1^W650A^* specifically in germ cells (using *nos-Gal4::VP16*) partially blocked differentiation, expression of *EcR-B1^F645A^* or *EcR-B1^DBL^* did not block egg chamber formation (Figure 2E-F,J). Expression of the *EcR-A* isoform or complementary versions of the dominant-negative *EcR-A* (*EcR-A^W622A^* and *EcR-A^F617A^*) recapitulated the *EcR-B1* phenotypes (Figure 2G-J). Since *EcR-B1^F645A^*/*EcR-A^F617A^*can bind ligand (while *EcR-B1^W6505A^*/*EcR-A^W622A^*blocks ligand binding), we postulate that they function as ligand sponges, effectively shunting endogenous EcR activity towards transcriptional repression. To further probe the effects of ligand concentration, we then tested whether high circulating ecdysone titer influenced the block to differentiation in *EcR-B1*-expressing females (Figure S2G-I). Feeding adult *EcR-B1* females 20-hydroxyecdysone for five days partially rescued germ cell differentiation. Taken together, these data suggest that the levels of EcR and ecdysone are tightly titrated in GSCs and cystoblasts to permit germ cell differentiation.

### Germline-specific over-expression of EcR stalls germline differentiation, phenocopying constitutive activation of BMP signaling

Germline-specific over-expression of *EcR* yielded a phenotype resembling germline manipulations of BMP signaling (Casanueva and Ferguson, 2004; McKearin and Spradling, 1990; Xie and Spradling, 1998). Like EcR, over-expression of a constitutively active form of the BMP receptor *tkv* (*UASp-tkv^ACT^*) or loss-of-function of the differentiation factor *bam* (*bam^RNAi^* or *bam^Δ86^*) resulted in germaria full of single and doublet cells (Figure 3A-D). Like wild-type ovaries, antibodies against phosphorylated Mad (pMad), a well-described read-out of active BMP signaling, highlighted cells closest to cap cells in *EcR-B1* germaria (Figure 3E-F). Similarly, cells lying closest to cap cells in *EcR-B1* ovaries expressed *Dad*, a transcriptional target of BMP signaling (Figure 3G-H). While germ cells in *EcR-B1* ovaries rarely expressed *bam::GFP*, a reporter of *bam* transcription (Figure 3I-J), most did not express Orb, an RNA binding protein enriched in oocytes in the final stages of mitotic division (Figure 3K-L) (Hinnant et al., 2020). In contrast, *EcR-B1*-expressing germ cells co-expressed Blanks, a nuclear protein that is silenced in differentiating germ cells (Figure 3M-N) (Kotb et al., 2024; Rust et al., 2020; Slaidina et al., 2021), and Chinmo, a transcription factor enriched in GSCs and undifferentiated germ cells (Figure 3O-P) (Grmai et al., 2024).

**Figure 3.**
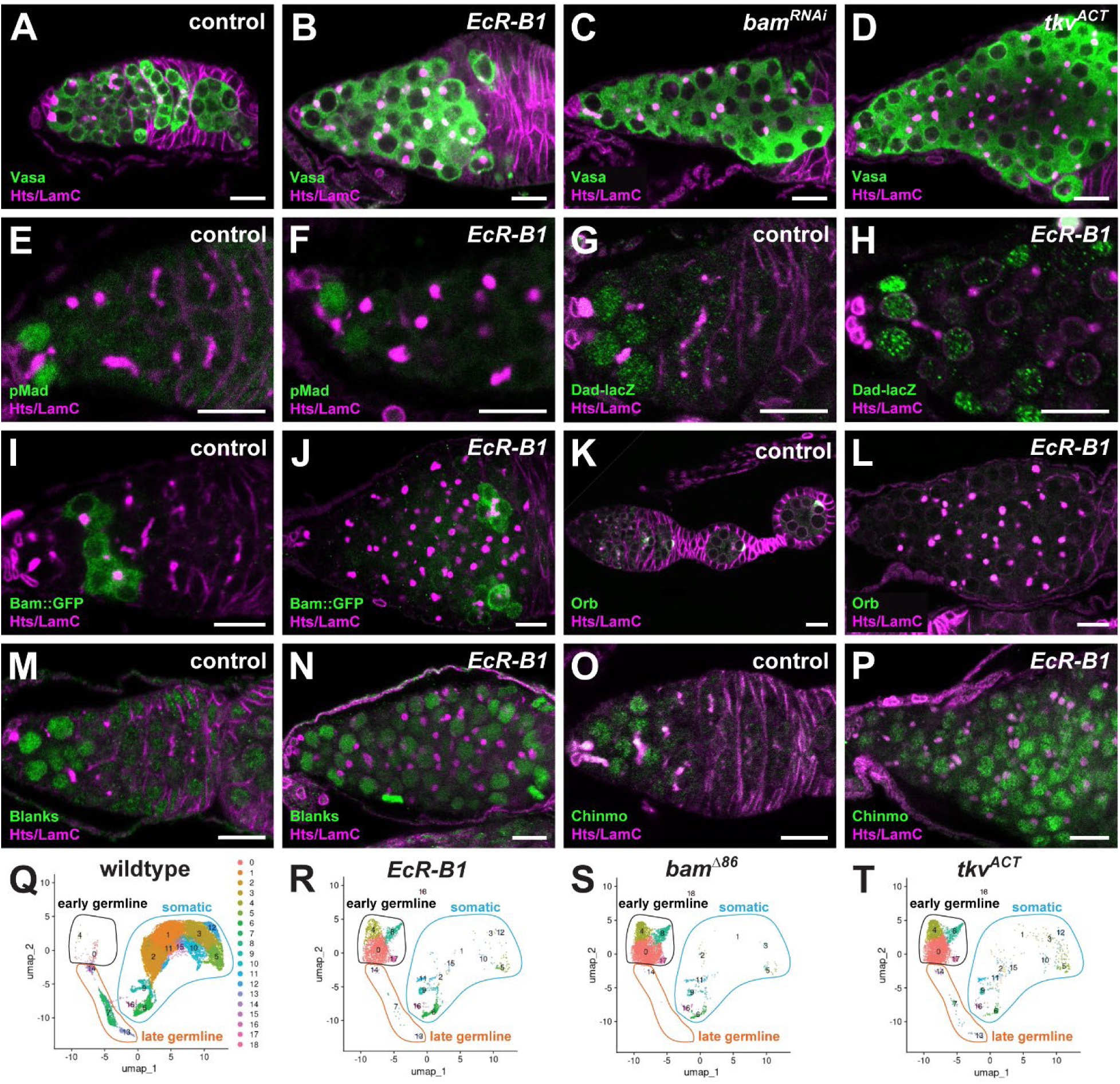
*EcR* over-expression phenocopies constitutive activation of BMP signaling. **(A-P)** *nos-Gal4::VP16* driving *UASp-lacZ* control (A,E,G,I,K,M,O), *UASz-EcR-B1* (B,F,H,J,L,N,P), *bam^RNAi^* (C), or *UASp-tkv^ACT^* (D) germaria immunostained for Hts and LamC (magenta; fusomes, follicle cells, and cap cells), DAPI (blue, nuclei), and Vasa (green, germ cells, A-D), pMad (green, active BMP signaling, E-F), β-galactosidase (green, Dad reporter, G-H), GFP (green, Bam reporter, I-J); Orb (green, oocytes, K-L), Blanks (green, undifferentiated germ cells, M-N), or Chinmo (green, undifferentiated germ cells, O-P). Scale bars = 10 µm. **(Q-T)** UMAP dimensionality reduction plots visualizing scRNA-seq analysis of all cell types from wild-type (Q, from (Slaidina et al., 2021), *nos-Gal4::VP16>UASz-EcR-B1* (R), *bam^ΔP^* (S), and *nos-Gal4::VP16>UASp-tkv^ACT^*(T) ovaries. Unique cell types (clusters) are numbered and displayed in differential colors, labeled for early germline (black outline), late germline (orange outline), or somatic cells (blue outline) based on differential gene expression (see Figure S3 for specific markers).

To assess similarities in overall cellular composition between the genotypes, we used single-cell RNA sequencing (scRNA-seq) to compare ovaries over-expressing *EcR-B1* or *tkv^ACT^* to wild-type ovaries (Slaidina et al., 2021) and ovaries harboring homozygous null mutation in *bam* (*bam^Δ86^*) (Sun et al., 2025). Germ cells were identified by enrichment for *nanos* and *vasa* and separated into undifferentiated and differentiated cells by expression of *Heterochromatin protein 6* and *minidiscs*, respectively (Figure S3) (Rust et al., 2020; Slaidina et al., 2021). Wild-type ovaries were primarily composed of somatic cells and contained few undifferentiated germ cells (Figure 3Q). In contrast, *EcR-B1*, *bam^Δ86^*, and *tkv^ACT^* ovaries were enriched for undifferentiated germ cells and depleted of both differentiated germ cells and most somatic cells, consistent with the reduced size of the tissue (Figure 3R-T). Taken together, these data indicate that over-expression of *EcR* in undifferentiated germ cells delays their differentiation prior to the mitotic-to-meiotic transition, phenocopying constitutively active *tkv* and *bam* loss of function.

### *EcR-B1* over-expression promotes *tkv* expression but does not block germ cell differentiation in a *tkv*- or *dpp*-dependent manner

To investigate transcriptional differences in germ cells over-expressing *EcR-B1* versus the other models, we used the scRNA-seq datasets to compare undifferentiated germ cells independently of the other ovarian cell types. Re-clustering of undifferentiated germ cells in wild-type, *bam^Δ86^*, *EcR-B1*, and *tkv^ACT^* germaria resulted in six subtypes (Figure 4A-E). Subtypes 0-3 included cells in wild-type ovaries, while subtypes 4 and 5 were unique, either to *bam^Δ86^* alone (subtype 4; Figure 4B) or all three genetic models (subtype 5; Figures 4B-D). Although undifferentiated cells across the four genotypes have transcriptional profiles similar enough to cluster into subtypes, cells in most clusters shared higher similarity to cells of the same genotype; that is, undifferentiated cells in subtypes 0 and 1 clustered in four independent groups, based on the genotype of the cells (Figure 4E). This is likely due to the differences in mRNA expression resulting from the genetic model itself. For example, *bam* mRNA was enriched across all clusters in *bam^Δ86^* cells (Figure 4F), as the *bam^Δ86^* mutation blocks protein production but not mRNA production(McKearin and Spradling, 1990). Similarly, *EcR* mRNA was specifically enriched in *UASz-EcR-B1*-expressing cells (Figure 4G).

**Figure 4.**
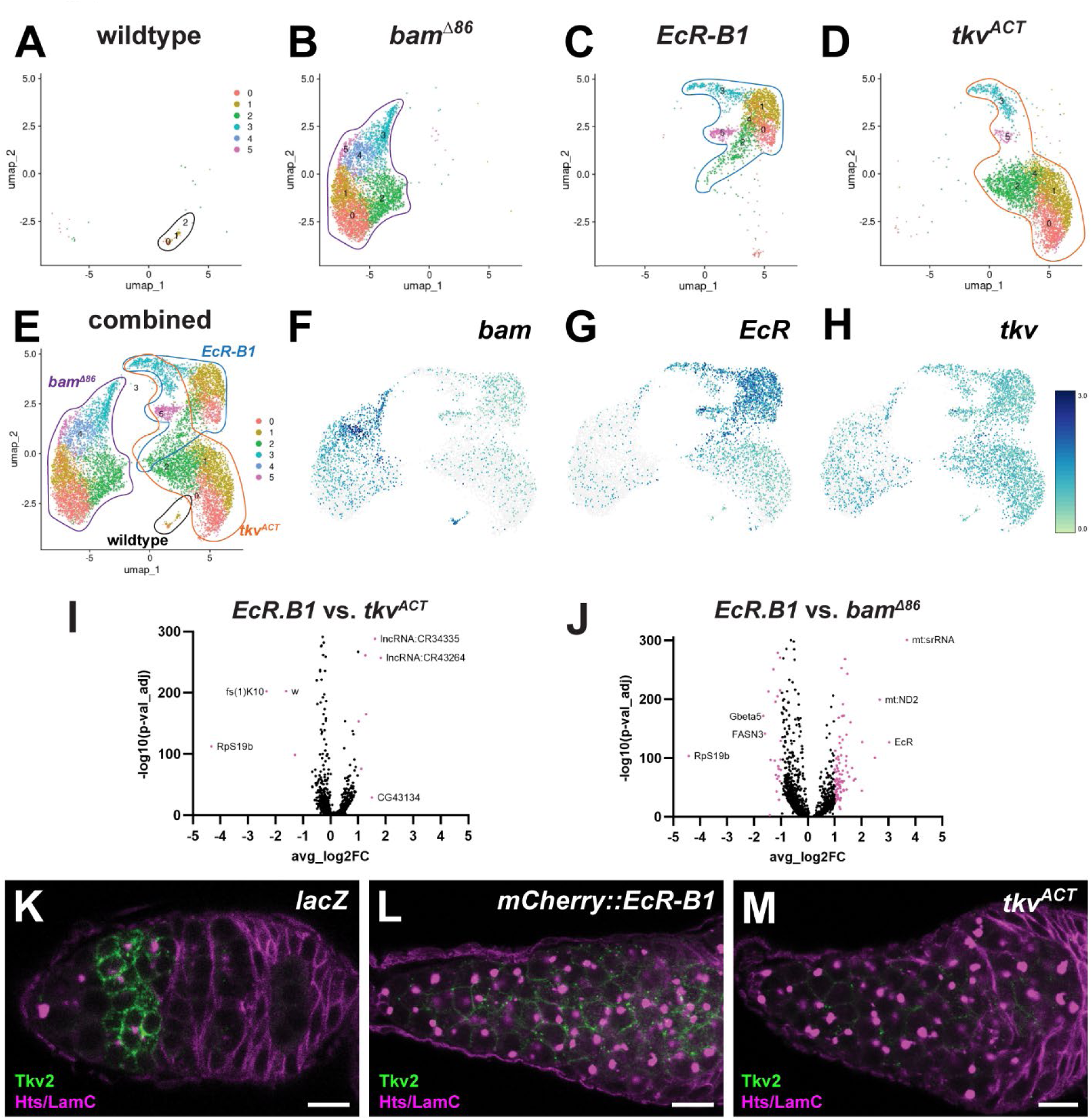
*EcR* over-expression promotes up-regulation of *tkv* in undifferentiated germ cells. **(A-E)** UMAP plots of undifferentiated germ cell sub-types from wild-type (A, from (Slaidina et al., 2021), *bam^Δ86^* (B), *nos-Gal4::VP16>UASz-EcR-B1* (C), and *nos-Gal4::VP16>UASp-tkv^ACT^* (D) ovaries. Six unique undifferentiated sub-types are displayed in differential colors (0, salmon; 1, gold; 2, green; 3, teal; 4, blue; 5, magenta). A single UMAP of undifferentiated germ cell sub-types merged from all four genotypes is displayed in (E), outlined for the individual genotypes (wild-type, black; *bam^Δ86^*, purple; *EcR-B1*, blue; *tkv^ACT^*, orange). **(F-H)** UMAP plots of undifferentiated germ cell subtypes overlayed with expression levels of *bam* (F), *EcR* (G), or *tkv* (H). Darker colors represent higher expression. **(I-J)** Volcano plots comparing differentially expressed genes in undifferentiated germ cell subtype 1 between *EcR-B1* and *tkv^ACT^* (I) and between *EcR-B1* and *bam^Δ86^*(J). **(K-M)** *nos-Gal4::VP16* driving *UASp-lacZ* control (K), *UASz-mCherry::EcR.B1* (L), or *UASp-tkv^ACT^* (M) germaria immunostained for Tkv (green) and Hts and LamC (magenta; fusomes, follicle cells, and cap cells). Scale bars = 10 µm.

Intriguingly, cells over-expressing *EcR-B1* or *tkv^ACT^*were highly similar, clustering together in overlapping profiles (Figure 4C-D). This suggested that over-expression of *EcR-B1* or *tkv^ACT^* resulted in similar gene regulation in undifferentiated cells. Indeed, while we expected that expression of *tkv* would be highest in *tkv^ACT^* -expressing cells, we found that *tkv* expression was up-regulated in response to both *EcR-B1* and *tkv^ACT^* over-expression (Figure 4H). Comparison of gene expression in individual subtypes revealed similar results. For example, few genes were differentially expressed in cells in subtype 1 between *EcR-B1* and *tkv^ACT^* germaria (Figure 4I), while more genes were differentially expressed between *EcR-B1* and *bam^Δ86^*germaria (Figure 4J).

To validate up-regulation of *tkv* in *EcR-B1* over-expressing cells *in situ*, we visualized Tkv protein levels in fixed tissue (Figure 4K-M) (Peterson et al., 2022). Although wild-type GSCs express Tkv, activated Tkv is degraded (Liu et al., 2023). This results in relatively low levels of Tkv protein until the 8-cell cyst stage, where it localized at germ cell plasma membranes (Figure 4K). Accordingly, *tkv^ACT^* germaria displayed low levels of Tkv protein in undifferentiated germ cells, like wild-type GSCs (Figure 4M). In contrast, Tkv protein was readily observed in *EcR-B1*-expressing germ cells away from the niche (Figure 4L), suggesting that *EcR-B1* over-expression specifically increases the levels of inactive Tkv protein. Intriguingly, however, the block to germ cell differentiation in *EcR*-expressing ovaries was not dependent on *dpp* or *tkv*, as loss of function of either gene failed to rescue germ cell differentiation (Figure S4). Thus, while *tkv* may be regulated by EcR, it is unlikely to be the sole factor impacting germ cell differentiation.

### Undifferentiated germ cells from *EcR-B1*, *tkv^ACT^*, and *bam^Δ86^* ovaries are a mixed population of mitotic cells

While the undifferentiated germ cell subtypes 0 and 1 diverged based on genetic background, cells in subtypes 2 and 3 clustered together independent of genotype, suggesting that over-expression of *EcR-B1*, constitutive activation of *tkv*, or loss of *bam* yield cells which converge at a similar cell type (Figure 4E). Consistent with this interpretation, computational temporal lineage analyses of the undifferentiated cell clusters based on pseudotemporal ordering or RNA kinetics (Monocle3 and scVelo, respectively) predict that subtypes 0 and 1 are most likely to be progenitors that differentiate into subtypes 2 and 3 (Figure S5).

Cells in subtype 1 are enriched with genes known to be expressed in GSCs (*CG2887*, *CG9705*, *CG32814*/*eggplant*, and *CG662836*; (Rust et al., 2020; Slaidina et al., 2021; Sun et al., 2023; Sun et al., 2025) and to regulate GSC maintenance, such as the cell cycle factors *CycB* and *CycE* (Ables and Drummond-Barbosa, 2013; Wang and Lin, 2005) and the asymmetrically distributed snoRNP component *wicked* (Fichelson et al., 2009). Gene ontology analysis of subtype 1 indicates enrichment of genes associated with “protein localization” and “RNA splicing”, consistent with GSCs. Further, cell cycle analysis of the undifferentiated subtypes demonstrates that while all undifferentiated cells are proliferative, cells in subtype 1 are less likely to be found in G1 phase of the cell cycle (Figure S6), consistent with the short G1 length displayed by GSCs (Ables and Drummond-Barbosa, 2013; Hinnant et al., 2017). In contrast, subtype 0 does not express high levels of cell cycle-related genes and instead is associated with ontologies “ATP synthesis”, “protein folding”, and “ribosome biogenesis”, associated with GSC differentiation (Sanchez et al., 2016; Teixeira et al., 2015). To corroborate the transcriptomics data, we identified three genes that are enriched in subtypes 0 and 1 and visualized their expression (Figure 5A-C’). The RNA helicase encoded by *Mov10* is enriched in subtype 1, consistent with endogenous expression in GSCs and pre-cystoblasts (Figure 5A-A’) (Takemura et al., 2021). Similarly, levels of the metabolic enzyme encoded by *ScsβA* and the chromosome-associated protein encoded by *HmgD* were similar across the undifferentiated subtypes, including subtypes 0 and 1. While a fluorescent reporter of *HmgD* is expressed throughout the germline (with lower levels in GSCs in G2 of the cell cycle), an *ScsβA* reporter is enriched in GSCs and cystoblasts, consistent with additional post-transcriptional regulation (Figure 5B-C’) (Buszczak et al., 2007). We conclude that subtypes 0 and 1 resemble endogenous GSCs/pre-cystoblasts.

**Figure 5.**
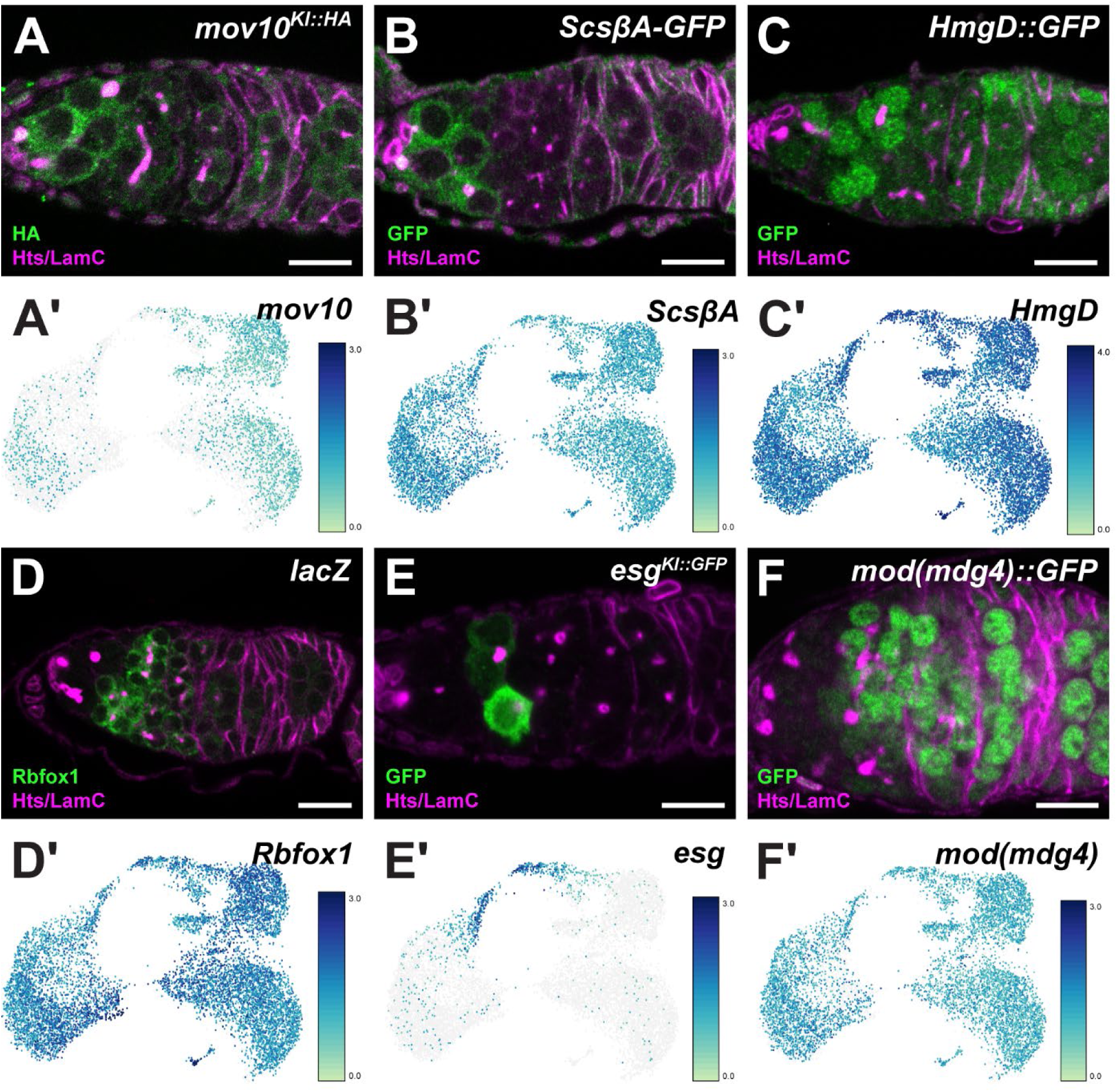
Undifferentiated germ cells from *EcR-B1*, *tkv^ACT^*, and *bam^Δ86^* ovaries are a mixed population primarily composed of GSCs, pre-cystoblasts, cystoblasts, and early dividing cysts. **(A-E)** Representative images (A-E) and expression levels (A’-E’) in undifferentiated germ cells from all four genotypes in the merged UMAP, visualizing *Mov10* (A-A’), ScsβA (B-B’), HmgD (C-C’), Rbfox1 (D-D’), *esg* (E-E’), and mod(mdg4) (F-F’). Germaria were immunostained for Hts and LamC (magenta; fusomes, follicle cells, and cap cells) and HA (A), GFP (B-C,E-F), or Rbfox1 (D) (green). Scale bars = 10 µm.

Germ cells in subtypes 2 and 3 were enriched for genes expressed in dividing cystoblasts/cysts, including the translation regulator *Rbfox* (Figure 5D-D’) (Carreira-Rosario et al., 2016), and associated with gene ontology terms “cytoplasmic translation”, “structural constituent of ribosome”, “transcription factor binding”, and protein kinase activity”, which are hallmarks of the transition from stem cell to differentiated germ cell (Breznak et al., 2023). Surprisingly, gene expression was largely non-overlapping between subtypes 2 and 3. We identified examples of genes uniquely expressed in subtype 3, including the transcription factor encoded by *esg*, which is transiently expressed in cystoblasts (Figure 5E-E’), despite its association with stem cell maintenance in other tissues (He et al., 2018). Many genes enriched in subtypes 2 and 3 are also highly expressed in differentiated germ cells. For example, expression of the chromatin regulator encoded by *mod(mdg4)* was equivalent across undifferentiated subtypes at the mRNA level; however, protein expression was suppressed in GSCs and enriched in differentiated cells (Figure 5F-F’). Similarly, genes typically associated with differentiated germ cells, such as *Nlg2*, *mnd*, and *slif*, were expressed in subtypes 2 and 3 (Rust et al., 2020; Slaidina et al., 2021). Consistent with prior studies, these results suggest that some maternal genes are transcriptionally activated during the first cyst division, either to regulate cyst growth or to be deposited into the early embryo (Pang et al., 2023; Samuels et al., 2024).

Although germ cells in *bam* loss-of-function and constitutively active BMP genetic backgrounds have been used extensively as models for GSC-like cells, our comparative analyses raised two possibilities. First, all three genetic models could stall differentiation, rather than block it completely, resulting in a mixed population of cells at non-equivalent differentiation states. Alternatively, the genetic perturbations could result in a population of cells stuck in a differentiation transition state, expressing markers of both stem cells and differentiated cells. To reconcile these possibilities, we again used Monocle3 pseudotime computational analysis to track the lineage of cells in each genotype (Figure S5D-F). While the trajectory of *EcR-B1* and *tkv^ACT^* germ cells ends in subtype 3, no endpoint could be identified in *bam^Δ86^* mutant germ cells. Instead, *bam^Δ86^* mutant germ cells exhibited a circuitous differentiation path, reverting to subtypes 0 and 1. These analyses suggest that *EcR-B1* and *tkv^ACT^*over-expression stall differentiation, while loss of *bam* pushes cells towards pre-cystoblast/cystoblast fates.

To further assess the differentiation state of *EcR-B1* germ cells, we visualized the GSC/cystoblast-enriched Mov10::HA and cyst-enriched Rbfox in *EcR-B1* germaria (Figure S7). As with Blanks and Chinmo (Figure 3), Mov10::HA expression expanded past the stem cell niche in *EcR-B1* germaria (Figure S7). Conversely, cytoplasmic Rbfox protein was largely absent from nearly all single and doublet germ cells, but could be easily visualized in the few cells that progressed into 4-cell cysts. We conclude that *bam* loss-of-function, BMP constitutive activation, and *EcR* over-expression result in discrete populations of GSC-like cells (closest to cap cells), pre-cystoblast-like cells (anterior germaria), and early dividing cystocytes (posterior germaria) caught between the stem cell and maternal transcriptional programs.

### A subset of canonical ecdysone-responsive genes is endogenously expressed in undifferentiated germ cells, suggesting a cell type-specific hormone response

EcR promotes cell type-specific transcriptional responses in target cells in part by activating a canonical transcriptional cascade of other nuclear receptors, chromatin binding proteins, and transcription factors (Uyehara et al., 2022; Uyehara and McKay, 2019). To elucidate direct and indirect transcriptional targets of EcR in undifferentiated germ cells, we queried whether canonical ecdysone-responsive genes were differentially expressed in *EcR-B1*, *tkv^ACT^*, or *bam^Δ86^* cells (Figure 6). We identified a subset of nuclear receptors and ecdysone-inducible genes up-regulated in *EcR-B1* and *tkv^ACT^* cells (Figure 6A). While prior studies demonstrated that *ftz-f1* is endogenously expressed in GSCs and dividing cysts and *Eip75B* is up-regulated in 16-cell cysts, consistent with its enrichment in subtype 3 (Beachum et al., 2021; Buszczak et al., 1999), we confirmed that *Hr78*, *Hnf4*, *Eip63E*, *E(bx)*, and *crol* are also endogenously expressed in germ cells (Figure 6B-F). Among these, *Eip75B* and *ftz-f1* are strong candidates as EcR direct targets. *EcR-B1*-expressing germ cells were enriched for *ftz-f1* and *Eip75B* expression, and *Eip75B* expression increased with differentiation in all three genetic backgrounds (Figure 6A). Depletion of *Eip75B* by germline-enhanced RNAi increased GSC number, while over-expression of *Eip75B* reduced GSC number (Figure S8), supporting a role as a repressor of GSC self-renewal (Ables and Drummond-Barbosa, 2010). In contrast to *EcR*, over-expression of *Eip75B* in germ cells did not block germ cell differentiation (Figure S8), suggesting that the increase in expression levels with stage of differentiation in the three differentiation-deficient genetic backgrounds reflects endogenous regulation of *Eip75B*.

**Figure 6.**
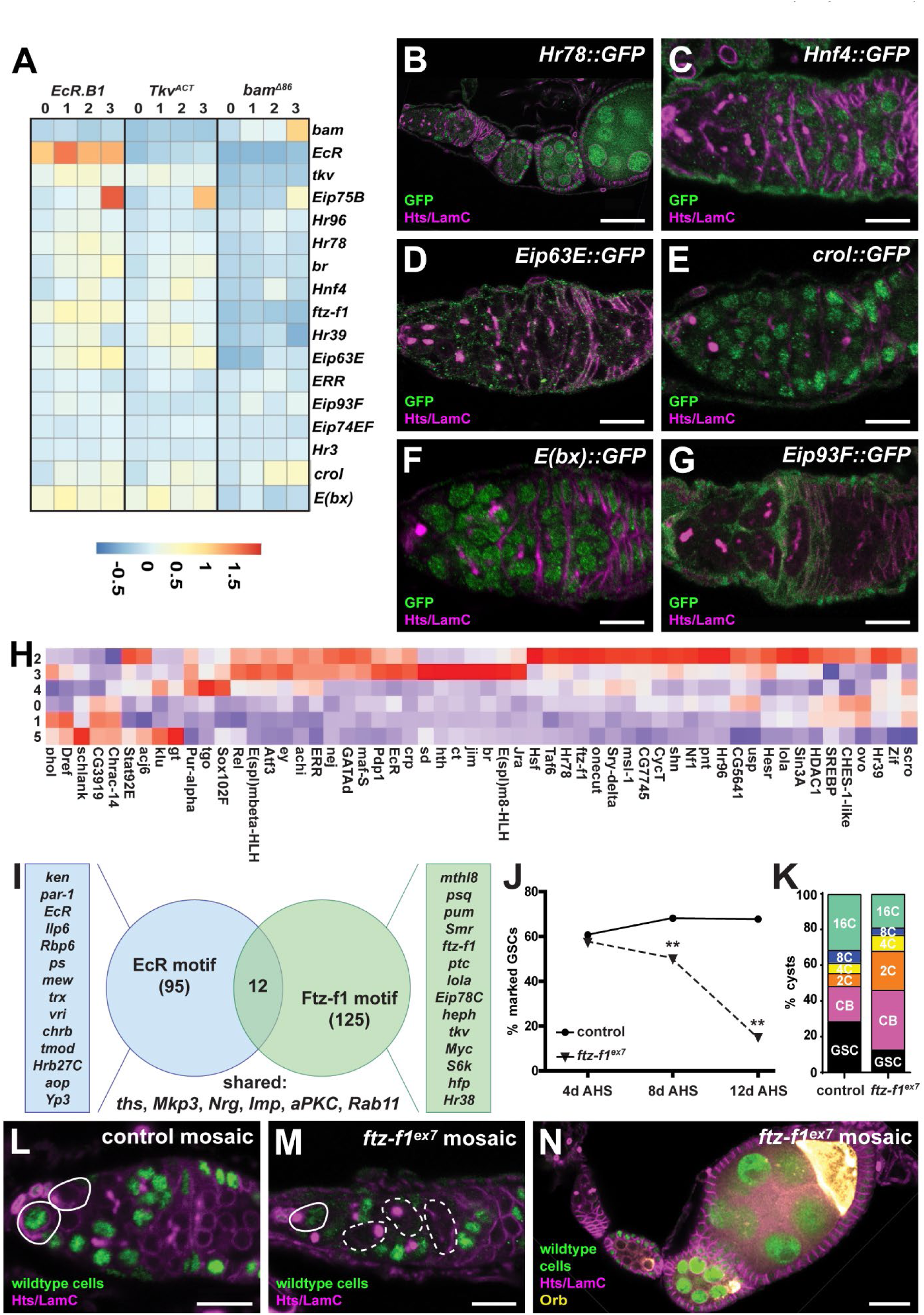
EcR elicits a cell type-specific transcriptional response in germ cells. **(A)** Heat map comparing expression of known ecdysone-responsive genes in *nos-Gal4::VP16>UASz-EcR-B1*, *nos-Gal4::VP16>UASp-tkv^ACT^*, and *bam^Δ86^* undifferentiated germ cells (subtypes 0-3). **(B-G)** Representative germaria from GFP-tagged reporter lines for *Hr78* (B), *HNF4* (C), *Eip63E* (D), *crol* (E), *E(bx)* (F), and *Eip93F* (G) immunostained for Hts and LamC (magenta; fusomes, follicle cells, and cap cells) and GFP (green, gene reporters). **(H)** Heat map showing transcription factor regulon enrichment in *bam^Δ86^* undifferentiated germ cells (all subtypes). **(I)** Comparison of putative EcR (blue) and Ftz-f1 (green) target genes in *bam^Δ86^* undifferentiated germ cells. Numbers represent total number of differentially-expressed genes containing a transcription factor binding motif. **(J-N)** Lineage tracing using *flippase*-inducible mosaic analysis compares the fate of control (*FRT79D*) and *ftz-f1^ex7^* null mutant germ cells. GSC self-renewal (J) was measured in germline mosaic germaria from control (circles) and *ftz-f1^ex7^* mutants (triangles) as the percentage of total germaria with a GFP-negative GSC at 4, 8, and 12 days after clone induction (n = at least 50 germaria per timepoint; \*\**p*<0.001, Chi-square test). Cyst distribution (K) compared the number of marked GSCs, cystoblasts (CB), or cysts (2C, 4C, 8C, and 16C) in mock mosaic control (left) or *ftz-f1^ex7^* mutant (right) germaria. Panels L-N show representative images from mock mosaic controls (L; where some cells are lineage labeled by the absence of GFP but all cells are wild-type) and *ftz-f1^ex7^* mutant ovarioles (M-N; where GFP-negative cells are homozygous null for *ftz-f1*) immunostained for GFP (green, wild-type cells), Hts and LamC (magenta), and Orb (yellow; panel N only). White outlines denote wild-type cells; dashed outlines denote *ftz-f1^ex7^* mutant cells). Scale bars = 10 µm (except B and N, = 20 µm).

Other known targets of EcR signaling, such as *ERR*, *Eip93F*, *Eip74EF*, and *Hr3*, were not differentially expressed between genotypes, suggesting that their regulation may be independent of EcR. Indeed, *Eip93F*, which is essential for ecdysone-dependent transcription in other cell types (Uyehara et al., 2017), is expressed exclusively in ovarian somatic cells (Figure 6G) and thus unlikely to mediate ecdysone-responsive transcription autonomously in germ cells. Notably, ecdysone biosynthesis gene expression was also absent from undifferentiated germ cells and instead enriched in somatic cells of the germarium (Figure S3), supporting the model that escort cells provide a local source of ecdysone ligands (Shi et al., 2021).

Since *bam^Δ86^* cells appear to block, rather than delay, differentiation at the cystoblast stage, we used this dataset to further identify gene regulatory networks active at this transition independently of our EcR genetic manipulation. Using Single Cell rEgulatory Network Interference and Clustering (SCENIC) analysis (Aibar et al., 2017) to predict potential gene regulatory networks active specifically in *bam^Δ86^* cells, we noted that cells in subtypes 2 and 3 were enriched in transcripts regulated by EcR, the obligate co-receptor Usp, or EcR-dependent transcription factors (Hr39, Ftz-f1, Hr78, Hr96, and Br; Figure 6H). As one example, we used DNA binding motif enrichment analysis to ask which genes regulated by EcR could also be regulated by Ftz-f1 (Figure 6I). Among 220 total enriched genes, only 12 contained both EcR and Ftz-f1 consensus binding sites, suggesting cascading regulation rather than co-regulation. Importantly, putative Ftz-f1 target genes included the transcriptional co-repressor encoded by *Smrter* (*Smr*), which is enriched in undifferentiated germ cells (Figure S9) and the nuclear receptor *Eip78C*, which is essential for cyst growth, packaging, and survival (Ables et al., 2015; Shi et al., 2021).

Taken together, gene regulatory network analyses suggest that EcR activity in undifferentiated cells regulates a transcription factor cascade during the earliest steps of germ cell differentiation. One prediction of this model is that an EcR-dependent transcription factor could regulate both GSC self-renewal and cyst differentiation. To test the model, we used clonal analysis to trace the lineage of *ftz-f1* null mutant germ cells (Figure 6J-N). We found that *ftz-f1* null mutant GSCs did not stay associated with the GSC niche over time, resulting in germaria lacking *ftz-f1* mutant GSCs but filled with single or doublet mutant germ cells (Figure 6J, L-M). While *ftz-f1* mutant germ cells could differentiate into 16-cell cysts (Figure 6N), their mitotic progression was greatly slowed, resulting in more mutant undifferentiated cells per germarium than controls (Figure 6K). We conclude that *ftz-f1* is essential for GSC self-renewal and germ cell differentiation and may regulate gene expression downstream of EcR.

### The EcR-dependent and BMP-dependent transcriptional programs converge on a transcriptional program regulating differentiation and cell cycle progression

The overlap in transcriptional repertoire between *EcR-B1*, *tkv^ACT^*, and *bam^Δ86^* undifferentiated germ cells suggested the possibility that EcR and BMP signaling might converge on similar transcriptional targets in GSCs and dividing cysts. To identify potential regulatory programs downstream of EcR and BMP activation, we combined all three scRNA-seq undifferentiated germ cell datasets and asked whether differentially expressed genes were enriched for common DNA binding factor motifs using SCENIC analysis (Figure 7A). Importantly, the predicted network controlling subtypes 0 and 1 include known regulators of self-renewal in the germline (i.e., Dp, Ovo, Nejire, and Maf-S) (Benner et al., 2024; Boija et al., 2017; Dynlacht et al., 1994; Kirilly et al., 2011; Pang et al., 2023; Tan et al., 2018). Among these, Dp binds the transcription factor E2F to promote cell cycle progression and is expressed primarily in dividing germ cells (Figure 7B), Ovo is a master regulator of germ cell transcription, Nejire is a chromatin binding protein known to stabilize open, active chromatin, and Maf-S is a transcription factor essential for GSC self-renewal in the testis (Tan et al., 2018). In contrast, the subtype 3 network includes factors that are maternally inherited and necessary for embryogenesis (i.e., Pdm3, Nubbin, Distalless, and Luna/KLF6(Ng et al., 1995; Özel et al., 2022; Plavicki et al., 2016; Weber et al., 2014)). Thus, subtype 3 likely represents the earliest transcription of maternally-expressed genes during germ cell differentiation.

**Figure 7.**
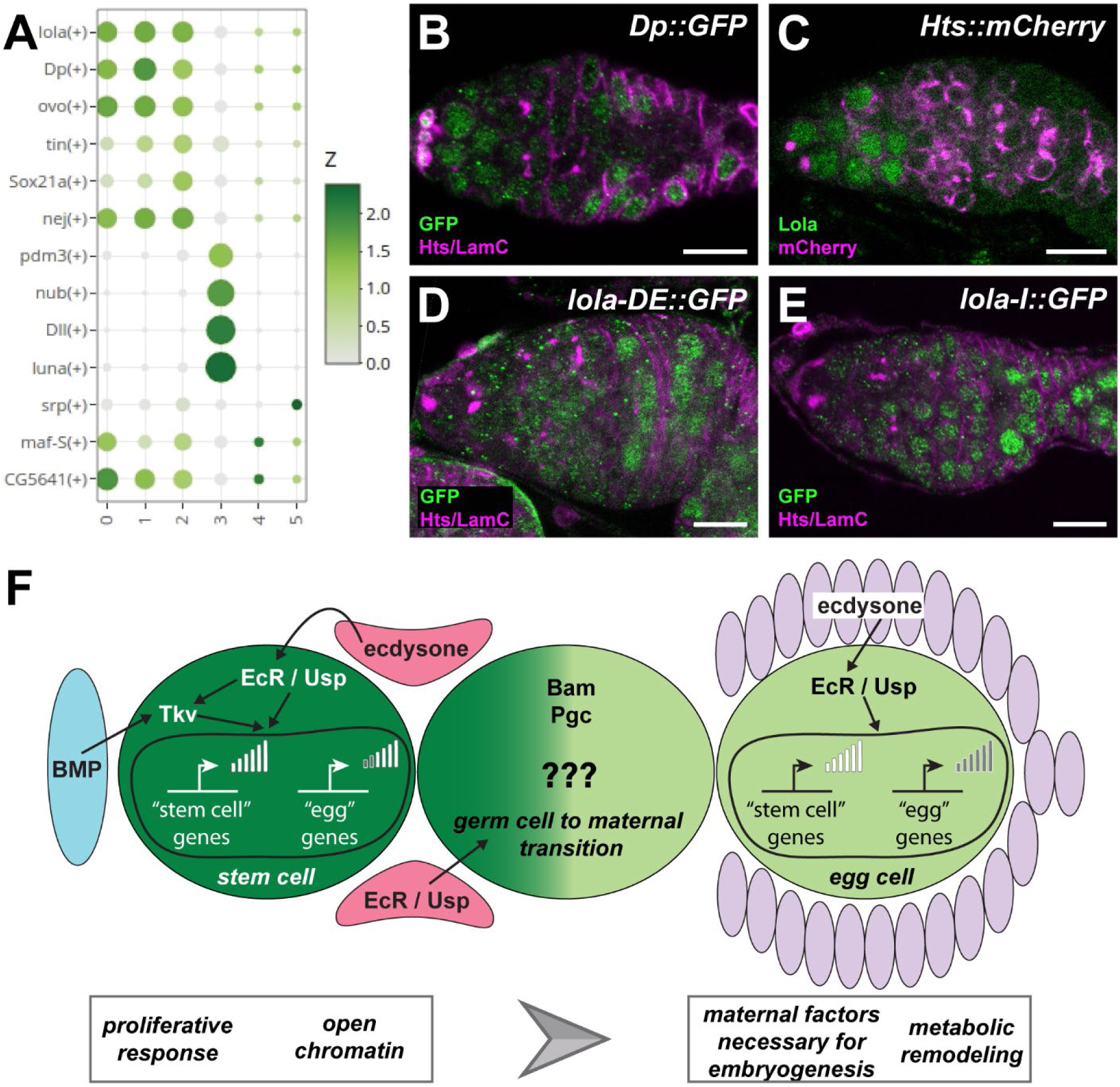
EcR and BMP signaling converge on a basal transcription program promoting cell cycle progression and differentiation. **(A)** Dot plot showing transcription factor regulon enrichment (regulon specificity score) in undifferentiated germ cells merged from all four genotypes. **(B-E)** Representative images of germaria from Dp (B) or Lola reporters (isoform *lola-DE*, D; isoform *lola-I*, E) immunostained for Hts and LamC (magenta; fusomes, follicle cells, and cap cells) and GFP (green, gene reporters), or wild-type germarium immunostained for Hts and LamC (magenta) and Lola (magenta). Scale bars = 10 µm. **(F)** Proposed model for EcR regulation of GSC self-renewal and germ cell differentiation. GSCs (pink) are maintained by EcR activation by low titer of ecdysone secreted by somatic cells (blue), which functions to promote GSC-self-renewal in part via a feed-forward loop promoting *tkv* expression. Independently, EcR activity in dividing cystoblasts/cysts promotes transcription of genes encoding maternal factors and necessary for metabolic remodeling, essential for reproductive success. This activity, combined with the non-autonomous action of EcR in adjacent cap and escort cells, collectively modulates GSC retention in the niche and germ cell differentiation.

SCENIC analysis also predicted regulation of germ cell differentiation by the transcription factor encoded by *longitudinals lacking* (*lola*; Figure 7A). The *lola* gene locus is complex, spanning over 61kb and known to encode at least 20 different proteins. Lola is required for gonad morphogenesis, stem cell maintenance, and differentiation (Bass et al., 2007; Davies et al., 2013; Silva et al., 2016; Zhao et al., 2022). SCENIC analysis predicted high Lola regulon activity in subtypes 0-2 (Figure 7A). To validate the computational analysis, we used an antibody against the Lola-DE isoform and available fluorescently-tagged transcriptional reporter lines to probe expression in wild-type germaria (Figure 7C-E). In contrast to pan-Lola antisera, which detect protein ubiquitously in the germarium (Zhao et al., 2022), anti-Lola-DE antisera detected a nuclear protein specifically enriched in GSCs and cystoblasts (Figure 7C). Intriguingly, however, C-terminal tagged fluorescent reporters of the *lola-DE* and *lola-I* isoforms were not expressed in undifferentiated cells and instead localized to differentiated 16-cell cyst nuclei (Figure 7D-E). These data suggests that Lola may influence gene expression programs during germ cell differentiation, and further highlight the complexity of gene regulation at the transcriptional and post-transcriptional level during the switch from stem cell to maternal factor expression. Notably, a *lola* paralog, *lola-like*, is essential for cyst growth (Kotb et al., 2025). Together, these analyses suggest that BMP and ecdysone signaling converge on a core program that controls fate transitions in the germline (Figure 7F).

## DISCUSSION

A key question in steroid hormone biology is how one hormone triggers distinct cellular responses, even within the same tissue. In *Drosophila*, ecdysone signals through the EcR, a heterodimeric receptor complex composed of two nuclear receptor proteins. Here, we demonstrate that although the EcR is expressed at roughly equivalent levels in neighboring cells in the *Drosophila* ovary, germ cells are controlled autonomously by EcR and respond differentially to EcR action depending on their state of differentiation. We propose that the unique chromatin status of GSCs and cystoblasts, combined with their unique spatial location in a BMP-enriched niche and exposure to low titer of ecdysone and EcR receptor, allow ecdysone signaling to elicit a unique transcriptional response in undifferentiated cells (Figure 7F). Together with non-autonomous EcR signaling in somatic cells, an EcR-dependent positive feedback loop may reinforce BMP signaling in GSCs by promoting expression of the BMP receptor *tkv*, while BMP signaling itself induces expression of an EcR-dependent transcriptional cascade. Finally, our data suggest that the higher levels of EcR in differentiated germ cells may stimulate a transcriptional cascade of additional nuclear receptors and other transcription factors that promote expression of genes necessary for cyst growth and germ cell maturation, as well as maternal mRNAs necessary for deposition into the developing oocyte. Given the broad functions of ecdysone signaling in the ovary, and the linear nature of differentiation, our data help establish the ovary as a powerful model to study how a steroid hormone elicits unique control over developing tissues.

### Defining an ecdysone-responsive transcriptional cascade in ovarian cells

Our current understanding of the ecdysone response in *Drosophila* cells was guided by the Ashburner model, a simplified framework for ligand-receptor interactions based on the observations of gene expression patterns in larval salivary glands (Morrow and Mirth, 2024; Thummel, 1996; Uyehara and McKay, 2019; Yamanaka et al., 2013). According to the model, low ecdysone titers induce expression of at least six “early” genes, including *Eip75B*, *br*, *Ftz-f1*, and *Eip93F,* thought to be direct targets of EcR. At high ecdysone titers, early gene products repress their own transcription and promote expression of a second set of more than 100 “late” genes, thought to be tissue-specific (Thummel, 1996). Nearly all the subsequent studies that refined and built upon this model analyzed the ecdysone response in somatic cells. This left open the possibility that ovarian germ cells, which have complex transcriptional and post-transcriptional mechanisms to regulate gene expression, could have unique molecular mechanisms for responding to hormone. Moreover, while the Ashburner model explains how a temporal gene expression cascade can be triggered by a single hormone pulse, it does not reconcile how individual cells make unique responses to the same hormone, nor how those responses are influenced by variables such as hormone and receptor titer and chromatin state.

Our data allow us to refine and build upon the Ashburner model in a tissue that is not dictated by a specific pulse of hormone, but rather reflects the steady state microenvironments within a hormone-producing tissue. In the adult ovary, it has long been established that late-stage ovarian cells (i.e. oogenesis stages 8-10, outside of the germarium) produce high titers of the hormone and express high levels of both EcR and Usp. In contrast, we confirm receptor expression is, in general, much lower in the germarium than in other cells, and identify escort cells of the germarium as a potential source of low-titer hormone production in the anterior ovary. Although we cannot definitively quantify the levels of each receptor isoform, our data suggest that receptor isoform is not a prevailing factor influencing unique cell responses to the hormone, but receptor titer likely governs the timing of cellular response. We also find that the differentiation status of ovarian cells is likely a defining factor for hormonal cell type specificity. Cells within the germarium microenvironment vary in developmental plasticity, including tissue-resident stem cells, differentiating cells, and terminally specified cells of multiple lineages. Accordingly, expression of the early response genes varies dependent on cell type, as *Eip75B* and *ftz-f1* predominate in GSCs and differentiating germ cells, while *br* and *Eip93F* are enriched in germarium somatic cells. Because chromatin status is a chief modifier of cellular plasticity and lineage, it is also likely to be a dominant regulator of the ecdysone response in these cells. GSCs/undifferentiated germ cells undergo two different chromatin states in quick succession. GSCs are rich in open euchromatin, while differentiating daughter cells are enriched for heterochromatin (Pang et al., 2023). Moreover, EcR is thought to preferentially bind open regions of chromatin, and ecdysone-response genes and/or EcR co-regulators may actively modulate chromatin state (Badenhorst et al., 2005; Carbonell et al., 2013; Kimura et al., 2008; Sedkov et al., 2003; Uyehara et al., 2022). Future studies comparing the chromatin state in individual cells of the germarium will provide further insight about how cells sense and respond in unique ways to the same hormone.

### Emerging questions about the control of hormone and receptor titer in the germline

The sensitivity of germ cells to ecdysone and EcR titer thus begs the question of what molecular mechanisms limit EcR activity/levels at the GSC/cystoblast/2-cell cyst transition. Our data raise several possibilities. First, EcR levels and/or activity may be kept low by a low titer of ecdysone. EcR regulates its own transcription and, in the wing disc, EcR/Usp functions as a transcriptional repressor at low ecdysone concentration (Morrow and Mirth, 2024; Nogueira Alves et al., 2022; Perez-Mockus et al., 2023). Similarly, since early germ cells do not appear to synthesize their own ecdysone (Figure S3) (Shi et al., 2021), we favor the model that anterior escort cells likely provide low levels of locally-produced ecdysone necessary for GSC maintenance and ligand-dependent EcR activity. As follicle cells from first wave egg chambers grow, both ovarian and circulating ecdysone concentration likely increase. Thus, EcR may repress itself in undifferentiated germ cells. Second, in contrast to the co-activator Taiman, which is neither abundant in undifferentiated germ cells nor required for GSC maintenance or cyst division (Ables and Drummond-Barbosa, 2010), the transcriptional co-repressor encoded by *Smrter* (analgous to mammalian *Nuclear Receptor Co-repressor 2*) is broadly expressed in GSCs, cystoblasts, and 2-cell cysts, but is down-regulated as cysts divide (Figure S9). The combination of higher ecdysone titer and lower co-repressor activity in 16-cell cysts may release the repression of EcR on itself.

A third possibility is that the sequential activation of ecdysone-dependent transcription factors may functionally limit EcR concentration and activity. Our data demonstrates up-regulation of the nuclear receptors *Eip75B*, *Eip78C*, and *ftz-f1* in dividing cysts (Figure 6), and suggests complex interaction between these proteins. We provide additional support for a role for *Eip75B* in limiting GSC self-renewal (Figure S8) (Ables and Drummond-Barbosa, 2010) and identify novel roles for f*tz-f1* in GSC self-renewal and cyst division (Figure 6). Both Eip75B and Ftz-f1 can repress *EcR* expression in other cellular contexts (Boulanger et al., 2011; Johnston et al., 2011). Moreover, Eip75B and Eip78C are paralogs (King-Jones and Thummel, 2005). *Eip78C* is weakly expressed in GSCs (our data and (Shi et al., 2021) and loss of *Eip78C* from adult GSCs does not impact their self-renewal (Ables et al., 2015); however, Eip78C and Eip75B may share overlapping transcriptional targets due to the similarity in their DNA binding domains. Since EcR and Ftz-f1 share few predicted transcriptional targets in cystoblasts (e.g. *bam^Δ86^*mutant cells), we conclude that EcR likely acts as self-limiting temporal switch for germ cell differentiation.

### The ovarian GSC microenvironment/niche as a unique model for studying the convergence of nuclear receptor signaling with other signaling pathways

The ability of stem cells to preserve tissue structure and optimize organ function relies on their ability to sense and respond to tissue-level and organism-level regenerative cues (Das et al., 2020; de Morree and Rando, 2023). A fundamental feature of this response is the organization of stem cells within tissue microenvironments (niches) that dictate their accessibility to local and systemic signals (Comazzetto et al., 2021; Xin et al., 2016). This is well-illustrated in the *Drosophila* ovary, where germline and follicle stem cells are centered at a nexus of diffusible signals (TGFβ/BMP, Wnt, EGFR, and Hh) that influence their fate (Losick et al., 2011). Yet how endocrine signals are received by stem cells in non-vascularized areas, and whether these cues are interpreted more specifically than would a general pro-proliferative or pro-survival signal, has been more difficult to decipher. Our study sheds light on this complex problem and offers an elegant model to understand hormonal regulation of stem cells in the context of the other signals that influence cellular activity. Consistent with previous work, we find that EcR modulates GSC self-renewal in part through a feedback loop with BMP signaling from adjacent somatic cells. We conclude that EcR and BMP signaling likely converge at the transcriptional level on a few key targets, including other transcription factors that module cell cycle progression and chromatin compaction. Our results highlight the remarkable cell-specific effects of steroid hormone signaling within a single tissue, illustrating how one hormone can drive dramatically different transcriptional outcomes in adjacent cells. Importantly, the GSC niche also likely protects cells in a relatively low-hormone environment. Although steroid hormones are typically considered long-range signals, we propose that local sources of steroid production or areas of receptor turnover or inactivation may likewise influence tissue-resident stem cell activity. Understanding how tissue microenvironments modify, shield, or concentrate steroid hormone titer or response may provide new avenues for modifying stem cell activity *in vivo*.

## EXPERIMENTAL PROCEDURES

### *Drosophila* strains and culture conditions

Flies were maintained at 22°-25°C on a standard medium containing cornmeal, molasses, yeast and agar (Nutrifly MF; Genesee Scientific). For all experiments, unless otherwise noted, flies were collected within 24 hours of eclosion and maintained at 25°C on standard medium supplemented with wet yeast paste for 5-6 days (changed daily) prior to ovary dissection. For *dpp* temperature sensitive mutants, crosses were set at 22°C and maintained at 18°C, progeny were collected within 24 hours of eclosion, and progeny were transferred to 25°C and fed for 5 days prior to dissection. Germline expression of transgenes or dsRNA-hairpins was facilitated by expressing the germline-specific *nos-GAL4::VP16-nos.UTR* (referred to throughout as *nos-Gal4::VP16*; Bloomington #4937) (Rørth, 1998; Van Doren et al., 1998). Driver expression was confirmed using *UASp-tubGFP* (Bloomington #7373) (Grieder et al., 2000) or UASp-lacZ (BDSC #98113) (Rørth, 1998). Females carrying *nos-Gal4::VP16* and *UASp-lacZ* were used as controls. A complete list of all transgenes used in this study can be found in Supplemental Table S1.

### Construction and validation of *UAS-EcR^RNAi^* transgenes

*UAS-EcR^RNAi^* lines were generated in *pVALIUM22* transgenes as described (www.flyrnai.org/TRiP-HOME.html) (Ni et al., 2011) for maximum germline efficiency. *UAS-EcR^RNAi2^*and *UAS-EcR^RNA3^* target an EcR coding exon common to all isoforms, using the same hairpin sequence as HMC03114 (CCGAGATGTGTTTCTCACTAA) and HMJ22371 (CTGCGATGCGAAGAAGAGCAA), respectively. *UAS-EcR^RNAi1^* was designed against the *EcR-A* isoform via the Designer of Small Interfering RNA tool (http://biodev.extra.cea.fr/DSIR/DSIRhtml) using default settings (21 nt siRNA; score threshold 90; CAACAGTAGCTACGCTAGATC); however, our analyses failed to confirm that the resulting dsRNA is specific to *EcR-A* (Supplemental Figure S1A-C). Primers were designed according to the Harvard Transgenic RNAi Project (TRiP) recommendations, annealed to form double-stranded oligos, and ligated into *EcoRI-HF*/*NheI–HF*–digested *pVALIUM22*. Transgenic flies were generated by phiC31 site-specific integrase into the attP2 site on the third chromosome (Bestgene). To validate *EcR* mRNA depletion in *UAS-EcR^RNAi^* lines, primers were designed to span the common coding region (exon 3 and 4) shared by *EcR-A* and *EcR-B1* isoforms, minimizing amplification of genomic DNA. All primers are listed in Supplemental Table S1.

To assess RNAi efficacy, germline knock-down was facilitated by *nos-GAL4::VP16* and RNA extraction was performed by dissecting 15 pairs of ovaries from females fed wet yeast paste for 5 days after eclosion. Ovaries were dissected in Invitrogen RNAlater (ThermoFisher/Invitrogen) solution and transferred to 1.5mL tubes containing RNAlater until time of extraction. Sterile technique was used by wiping all gloves, surfaces, and instruments with Invitrogen RNaseZap. Whole-ovary RNA extraction was performed using the RNAqueous-4PCR RNA Isolation kit (ThermoFisher/Invitrogen). RNA concentration and purity were determined using the Agilent Bioanalyzer RNA Pico assay while being kept at -80°C. Genetic replicate samples were used to generate cDNA using iScript synthesis kit (Bio-Rad). RT-PCR amplification of generated cDNA was run at 25 cycles at T_m_ = 57°C for three genetic replicates. PCR products were run on a 1.5% agarose gel to measure intensity levels. Levels were compared across replicate samples.

### Construction of *EcR* over-expression and dominant negative transgenes

To generate *UASz*-*EcR* over-expression and dominant-negative constructs, the *EcR* coding region with or without nucleotide substitutions was synthesized into *pUC57* mini (GenScript) following the methods of Cherbas and colleagues (Cherbas et al., 2003; Hu et al., 2003), creating *pUC57-EcR-B1^ΔC655^*, *pUC57-EcR-B1^ΔC655-F645A^*, *pUC57-EcR-B1^ΔC655-W650A^*, and *pUC57-EcR-B1^ΔC655-F645A+W650A^* (referred to as *pUC57-EcR-B1^DBL^*). Likewise, the coding region for *EcR-A* was designed with sequences equivalent to *EcR-B1* (*pUC57-EcR-A^ΔC627^*). Coding regions were amplified from *pUC57* mini and cloned into the *BamHI* site of *pUASz-1.0* (DeLuca and Spradling, 2018) or a modified *pUASz-1.0* with *mCherry* in frame at the C- or N-terminus using InFusion Snap Assembly Cloning (Takara). *EcR-A* dominant negative transgenes [*EcR-A^ΔC627-F617A^*, *EcR-A^ΔC627-W622A^*, and *EcR-A^ΔC627-F617A+W622A^* (referred to as *EcR-A^DBL^*)] were cloned by site directed mutagenesis in *pUASz-EcR-A*. Plasmid sequences were verified via nanopore sequencing (Plasmidsaurus). Plasmid DNA was injected into embryos by BestGene and docked into the *attP40* locus on chromosome two by phiC31-mediated integration.

### Genetic mosaic generation

For genetic mosaic analyses with negative labeling using flippase (FLP)/FLP recognition target (FRT), we used the null allele *ftz-f1^ex7^*(BDSC #64338) on *FRT79D*. Other genetic tools are described in FlyBase version FB 2025_04 (Öztürk-Çolak et al., 2024). Genetic mosaics were generated by FLP/FRT-mediated recombination in 2-3-day old females carrying a mutant allele in trans to a wildtype allele (linked to a *Ubi-nGFP* marker) on homologous FRT arms, and a *hs-FLP* transgene, as described (Laws and Drummond-Barbosa, 2015). For induction of transgene expression by FLP-out genetic recombination, we used *hsFLP; nos-FRT-STOP-FRT-Gal4, UASp-H4-eGFP* (a gift of T. Xie) (Phipps et al., 2023). Flies were heat shocked at 37°C two times per day for three days and incubated at 25°C supplemented with wet yeast paste prior to dissection. Wild-type alleles were used for the generation of control (mock) mosaics.

### Immunofluorescence and microscopy

Ovaries were dissected, fixed, washed, and blocked as previously described (Ables et al., 2016; Hinnant et al., 2017; Williams and Ables, 2023). In the standard protocol, ovaries were dissected and teased apart in Grace’s media (Caisson Labs) and fixed in cold 5.3% formaldehyde in Grace’s media at room temperature for 13 minutes. A modified protocol was also used for some antibodies, wherein ovaries were fixed with room temperature 5.8% formaldehyde in Grace’s Insect Medium for 10 minutes at room temperature. Ovaries were washed extensively in phosphate-buffered saline (PBS, pH 7.4; Thermo Fisher) with 0.1% Triton X-100 (PBST), permeabilized in PBS with 0.5% Triton X-100, then blocked for one to three hours in a blocking solution consisting of 5% bovine serum albumin (Sigma), 5% normal goat serum (MP Biomedicals) and PBST. Samples were incubated with primary antibodies in block at 4°C overnight. Samples were incubated with Alexa Fluor 488-, 568- or 633-conjugated goat-species specific secondary antibodies (ThermoFisher; 1:200). A complete list of antibodies is listed in Supplemental Table S1. Where appropriate, EdU was detected using AlexaFluor-594 or -647 via Click-It chemistry following the manufacturer’s recommendations (ThermoFisher). Ovaries were counterstained with DAPI (Sigma 1:1000 in PBS). Ovaries were then mounted in 90% glycerol containing 20.0 µg/mL N-propyl gallate (Sigma). Data was collected using a Zeiss LSM 700 laser scanning confocal microscope equipped with a 63X plan apo objective. Images were analyzed using Zen Black software and images were minimally and equally enhanced via histogram using Zen and Adobe Photoshop CC.

### Germ cell analyses and statistics

GSCs were identified based on the juxtaposition of their fusomes (at any morphology) to the junction with adjacent cap cells (Ables and Drummond-Barbosa, 2010; Villa-Fombuena et al., 2021). Germ cell differentiation was scored as the number of germaria with blocked differentiation phenotype, compared to controls and subjected to Fisher’s exact T-test analyses using Prism (GraphPad). In *FLP/FRT* mosaics, GSC loss was measured as the percentage of total germaria analyzed that had at least one GFP-negative GSC, and subjected to Fisher’s exact T-test (Laws and Drummond-Barbosa, 2015). In RNAi progeny, GSC loss was measured as the average number of GSCs present over three timepoints. Results were subjected to Student’s two-tailed T-test at each timepoint using Microsoft Excel and Prism.

**Single Cell RNA sequencing and analyses.**

Using Method A (Slaidina et al., 2021), 100 ovaries from each genotype were dissected in Grace’s insect medium (Caisson Labs) and disassociated in a solution containing 0.5% type 1 collagenase and 1% trypsin in PBS solution for 15 minutes using vigorous pipetting every few minutes. Disassociation was stopped by adding Grace’s insect medium with 10% fetal bovine serum. Suspended cells were strained through a Flowmi 40-micron filter tip (Sigma-Aldrich) and then centrifuged at 5 rpm for 3 minutes. Supernatant was decanted, cells were resuspended in 0.04% BSA in PBS and centrifuged again at 5 rpm for 3 minutes. Final suspension was in Grace’s insect medium and then preserved on ice. Live/dead count was measured on the Invitrogen Countess 3FL Cell Counter. 1.70×10^6^ cells from the *nos-Gal4>UASp-tkv^ACT^*sample were collected (89% viability) and 1.17×10^7^ cells from the *nos-Gal4>UASz-EcR-B1* sample were collected (95% viability).

Libraries were constructed using 10X Genomics Chromium single cell 3’ V3 kit according to the manufacturer’s protocol. Sequencing of the pooled libraries were performed on the NextSeq 2000 platform (Illumina). For the *EcR-B1* sample, 5,741 cells were sequenced with a total of 136,771,995 reads (an average of 23,799 reads per cell; 96.70% valid barcodes; 74.6% mapped confidently to the genome; 4,112 median UMI counts per cell). For the *tkv^ACT^* sample, 7,043 cells were sequenced with a total of 116,346,480 reads (an average of 16,519 per cell; 97.40% valid barcodes; 96.8% mapped confidently to the genome; 6,934 median UMI counts per cell). Data can be accessed from the NCBI GEO database (GSE309824).

Raw sequencing reads in FASTQ files were obtained using cellranger (v7.2.0) mkfastq. The alignment to *Drosophila melanogaster* reference genome BDGF6.46 (Ensembl) and the followed quantification was conducted using cellranger count. Wild-type ovary scRNA-seq raw sequencing data were downloaded from NCBI GEO database (GSE162192). *bam^Δ86^* raw sequencing data were provided ahead of publication and can be accessed at https://bigd.big.ac.cn/ (CRA028191), from which 6800 single cells were randomly selected for analysis (to keep the total number of cells comparable to those from *EcR-B1* and *tkv^ACT^* samples). Subsequently, Seurat (v5.1.0) package in RStudio (Build 369) with R (v4.4.1) were used for data analysis. Briefly, single cells were firstly filtered with nFeature_RNA (more than 1000 but less than 6000). Doublets or multiplets were identified by DoubletFinder (v2.0.4), mainly in the GC of control samples (4cc, 8cc, and 16cc) and were thus not eliminated. Batch effects were corrected using Harmony (v1.2.0). Gene expression data were then aggregated, normalized and variance stabilized with sctransform regularized negative binominal regression (SCTransform in Seurat). Dimensionality reduction was performed by principal component analysis (PCA) and uniform manifold approximation and projection (UMAP) embedding (dims = 1:30) with RunPCA and RunUMAP, respectively. Cell clusters were subsequently identified with FindNeighbors (dims = 1:30) and FindClusters (resolution = 0.2 for initial clustering and 0.5 for refined clustering); and annotated based on expression of cell type-specific gene biomarkers. Differentially expressed biomarkers among cell types were assessed with FindAllMarkers or FindMarkers for genes detected in at least 10% of cells using a log(foldchange) threshold at 0.25. Bonferroni-adjusted p-values were used to determine significance at a false discovery rate (FDR) < 0.05.

The aggregated data set of wild-type (GSE162192), *bam^Δ86^* (CRA028191), *EcR-B1* and *tkv^ACT^* (GSE309824) were visualized using the 10X Loupe Browser platform. The aggregated data of both samples pipeline-generated 16 clusters. When measuring differentially expressed genes between clusters and groups, Log2 fold change is the ratio of the normalized mean gene UMI counts in the cluster relative to all other clusters. P-values were adjusted using the Benjamini-Hochberg correction for multiple tests.

## AUTHOR CONTRIBUTIONS

Conceptualization: L.J., A.W., E.T.A.

Data curation: C.Y., W.H., Z.S., T.N., E.T.A.

Formal Analysis: L.J., A.W., C.Y., E.T.A.

Funding acquisition: E.T.A.

Investigation: L.J., A.W., C.Y., W.H., A.S., L.S., V.G., D.P., S.M., B.C., Z.S., E.T.A.

Methodology: L.J., E.T.A.

Project administration: E.T.A.

Resources: Z.S., T.N.

Software: C.Y., W.H., Z.S.

Supervision: L.J., A.W., E.T.A.

Validation: L.J., A.W., A.S., L.S., V.G., E.T.A.

Visualization: L.J., A.W., E.T.A.

Writing – original draft preparation: L.J., A.W., E.T.A.

Writing – review and editing: L.J., A.W., W.H., A.S., L.S., V.G., D.P., B.C., Z.S., T.N., E.T.A.

## ACKNOWLEDGMENTS

We are grateful to Daniela Drummond-Barbosa, Erik Sontheimer, Nick Sokol, Lesley Weaver, Dan McKay, Ting Xie, Paul Lasko, Hiroshi Nakato, Michael O’Connor, Michael Buszczak, Prashanth Rangan, the Developmental Studies Hybridoma Bank, the *Drosophila* Genetic Resource Center, the *Drosophila* Research & Screening Center-Biomedical Technology Research Resource at Harvard, and the Bloomington *Drosophila* Stock Center for antibodies and fly lines. Many thanks to Amanda Powell and Sophia Spohn for technical assistance, and to Lauren Anllo, Beth Thompson, Amanda Powell, Dan McKay, and Prashanth Rangan for helpful discussion and critical feedback on this work.

## REFERENCES

Ables, E. T., Bois, K. E., Garcia, C. A. and Drummond-Barbosa, D. (2015). Ecdysone response gene E78 controls ovarian germline stem cell niche formation and follicle survival in Drosophila. Dev Biol 400, 33–42.

Ables, E. T. and Drummond-Barbosa, D. (2010). The steroid hormone ecdysone functions with intrinsic chromatin remodeling factors to control female germline stem cells in Drosophila. Cell Stem Cell 7, 581–592.

Ables, E. T. and Drummond-Barbosa, D. (2013). Cyclin E controls Drosophila female germline stem cell maintenance independently of its role in proliferation by modulating responsiveness to niche signals. Development 140, 530–540.

Ables, E. T., Hwang, G. H., Finger, D. S., Hinnant, T. D. and Drummond-Barbosa, D. (2016). A Genetic Mosaic Screen Reveals Ecdysone-Responsive Genes Regulating Drosophila Oogenesis. G3 (Bethesda, Md.) 6, 2629–2642.

Aibar, S., González-Blas, C. B., Moerman, T., Huynh-Thu, V. A., Imrichova, H., Hulselmans, G., Rambow, F., Marine, J. C., Geurts, P., Aerts, J., et al. (2017). SCENIC: single-cell regulatory network inference and clustering. Nat Methods 14, 1083–1086.

Ameku, T. and Niwa, R. (2016). Mating-Induced Increase in Germline Stem Cells via the Neuroendocrine System in Female Drosophila. PLoS Genet 12, e1006123.

Badenhorst, P., Xiao, H., Cherbas, L., Kwon, S. Y., Voas, M., Rebay, I., Cherbas, P. and Wu, C. (2005). The Drosophila nucleosome remodeling factor NURF is required for Ecdysteroid signaling and metamorphosis. Genes Dev 19, 2540–2545.

Bass, B. P., Cullen, K. and McCall, K. (2007). The axon guidance gene lola is required for programmed cell death in the Drosophila ovary. Dev Biol 304, 771–785.

Beachum, A. N., Whitehead, K. M., McDonald, S. I., Phipps, D. N., Berghout, H. E. and Ables, E. T. (2021). Orphan nuclear receptor ftz-f1 (NR5A3) promotes egg chamber survival in the Drosophila ovary. G3 (Bethesda, Md.) 11.

Benner, L., Muron, S., Gomez, J. G. and Oliver, B. (2024). OVO positively regulates essential maternal pathways by binding near the transcriptional start sites in the Drosophila female germline. Elife 13.

BharathwajChetty, B., Sajeev, A., Vishwa, R., Aswani, B. S., Alqahtani, M. S., Abbas, M. and Kunnumakkara, A. B. (2024). Dynamic interplay of nuclear receptors in tumor cell plasticity and drug resistance: Shifting gears in malignant transformations and applications in cancer therapeutics. Cancer Metastasis Rev 43, 321–362.

Boija, A., Mahat, D. B., Zare, A., Holmqvist, P. H., Philip, P., Meyers, D. J., Cole, P. A., Lis, J. T., Stenberg, P. and Mannervik, M. (2017). CBP Regulates Recruitment and Release of Promoter-Proximal RNA Polymerase II. Mol Cell 68, 491–503.e495.

Boulanger, A., Clouet-Redt, C., Farge, M., Flandre, A., Guignard, T., Fernando, C., Juge, F. and Dura, J. M. (2011). ftz-f1 and Hr39 opposing roles on EcR expression during Drosophila mushroom body neuron remodeling. Nat Neurosci 14, 37–44.

Breznak, S. M., Kotb, N. M. and Rangan, P. (2023). Dynamic regulation of ribosome levels and translation during development. Seminars in cell & developmental biology 136, 27–37.

Buszczak, M., Freeman, M. R., Carlson, J. R., Bender, M., Cooley, L. and Segraves, W. A. (1999). Ecdysone response genes govern egg chamber development during mid-oogenesis in Drosophila. Development 126, 4581–4589.

Buszczak, M., Paterno, S., Lighthouse, D., Bachman, J., Planck, J., Owen, S., Skora, A. D., Nystul, T. G., Ohlstein, B., Allen, A., et al. (2007). The carnegie protein trap library: a versatile tool for Drosophila developmental studies. Genetics 175, 1505–1531.

Carbonell, A., Mazo, A., Serras, F. and Corominas, M. (2013). Ash2 acts as an ecdysone receptor coactivator by stabilizing the histone methyltransferase Trr. Mol Biol Cell 24, 361–372.

Carreira-Rosario, A., Bhargava, V., Hillebrand, J., Kollipara, R. K., Ramaswami, M. and Buszczak, M. (2016). Repression of Pumilio Protein Expression by Rbfox1 Promotes Germ Cell Differentiation. Developmental cell 36, 562–571.

Casanueva, M. O. and Ferguson, E. L. (2004). Germline stem cell number in the Drosophila ovary is regulated by redundant mechanisms that control Dpp signaling. Development 131, 1881–1890.

Cherbas, L., Hu, X., Zhimulev, I., Belyaeva, E. and Cherbas, P. (2003). EcR isoforms in Drosophila: testing tissue-specific requirements by targeted blockade and rescue. Development 130, 271–284.

Chhabra, S. N. and Booth, B. W. (2021). Asymmetric cell division of mammary stem cells. Cell Div 16, 5.

Comazzetto, S., Shen, B. and Morrison, S. J. (2021). Niches that regulate stem cells and hematopoiesis in adult bone marrow. Developmental cell 56, 1848–1860.

Das, D., Fletcher, R. B. and Ngai, J. (2020). Cellular mechanisms of epithelial stem cell self-renewal and differentiation during homeostasis and repair. Wiley Interdiscip Rev Dev Biol 9, e361.

Davies, E. L., Lim, J. G., Joo, W. J., Tam, C. H. and Fuller, M. T. (2013). The transcriptional regulator lola is required for stem cell maintenance and germ cell differentiation in the Drosophila testis. Dev Biol 373, 310–321.

de Morree, A. and Rando, T. A. (2023). Regulation of adult stem cell quiescence and its functions in the maintenance of tissue integrity. Nat Rev Mol Cell Biol 24, 334–354.

DeLuca, S. Z. and Spradling, A. C. (2018). Efficient Expression of Genes in the Drosophila Germline Using a UAS Promoter Free of Interference by Hsp70 piRNAs. Genetics 209, 381–387.

Drummond-Barbosa, D. (2019). Local and Physiological Control of Germline Stem Cell Lineages in Drosophila melanogaster. Genetics 213, 9–26.

Dynlacht, B. D., Brook, A., Dembski, M., Yenush, L. and Dyson, N. (1994). DNA-binding and trans-activation properties of Drosophila E2F and DP proteins. Proc Natl Acad Sci U S A 91, 6359–6363.

Fichelson, P., Moch, C., Ivanovitch, K., Martin, C., Sidor, C. M., Lepesant, J. A., Bellaiche, Y. and Huynh, J. R. (2009). Live-imaging of single stem cells within their niche reveals that a U3snoRNP component segregates asymmetrically and is required for self-renewal in Drosophila. Nat Cell Biol 11, 685–693.

Grieder, N. C., de Cuevas, M. and Spradling, A. C. (2000). The fusome organizes the microtubule network during oocyte differentiation in Drosophila. Development 127, 4253–4264.

Grmai, L., Jimenez, E., Baxter, E. and Doren, M. V. (2024). Steroid signaling controls sex-specific development in an invertebrate. bioRxiv.

Harshman, L. G., Loeb, A. M. and Johnson, B. A. (1999). Ecdysteroid titers in mated and unmated Drosophila melanogaster females. J Insect Physiol 45, 571–577.

He, L., Si, G., Huang, J., Samuel, A. D. T. and Perrimon, N. (2018). Mechanical regulation of stem-cell differentiation by the stretch-activated Piezo channel. Nature 555, 103–106.

Hinnant, T. D., Alvarez, A. A. and Ables, E. T. (2017). Temporal remodeling of the cell cycle accompanies differentiation in the Drosophila germline. Dev Biol 429, 118–131.

Hinnant, T. D., Merkle, J. A. and Ables, E. T. (2020). Coordinating Proliferation, Polarity, and Cell Fate in the Drosophila Female Germline. Front Cell Dev Biol 8, 19.

Hodgetts, R. B., Sage, B. and O’Connor, J. D. (1977). Ecdysone titers during postembryonic development of Drosophila melanogaster. Dev Biol 60, 310–317.

Hsu, Y. C. and Fuchs, E. (2022). Building and Maintaining the Skin. Cold Spring Harb Perspect Biol 14.

Hu, X., Cherbas, L. and Cherbas, P. (2003). Transcription activation by the ecdysone receptor (EcR/USP): identification of activation functions. Mol Endocrinol 17, 716–731.

Johnston, D. M., Sedkov, Y., Petruk, S., Riley, K. M., Fujioka, M., Jaynes, J. B. and Mazo, A. (2011). Ecdysone- and NO-mediated gene regulation by competing EcR/Usp and E75A nuclear receptors during Drosophila development. Mol Cell 44, 51–61.

Kimura, S., Sawatsubashi, S., Ito, S., Kouzmenko, A., Suzuki, E., Zhao, Y., Yamagata, K., Tanabe, M., Ueda, T., Fujiyama, S., et al. (2008). Drosophila arginine methyltransferase 1 (DART1) is an ecdysone receptor co-repressor. Biochem Biophys Res Commun 371, 889–893.

King-Jones, K. and Thummel, C. S. (2005). Nuclear receptors--a perspective from Drosophila. Nat Rev Genet 6, 311–323.

Kirilly, D., Wong, J. J., Lim, E. K., Wang, Y., Zhang, H., Wang, C., Liao, Q., Wang, H., Liou, Y. C., Wang, H., et al. (2011). Intrinsic epigenetic factors cooperate with the steroid hormone ecdysone to govern dendrite pruning in Drosophila. Neuron 72, 86–100.

König, A. and Shcherbata, H. R. (2015). Soma influences GSC progeny differentiation via the cell adhesion-mediated steroid-let-7-Wingless signaling cascade that regulates chromatin dynamics. Biol Open 4, 285–300.

König, A., Yatsenko, A. S., Weiss, M. and Shcherbata, H. R. (2011). Ecdysteroids affect Drosophila ovarian stem cell niche formation and early germline differentiation. Embo j 30, 1549–1562.

Kotb, N. M., Ulukaya, G., Chavan, A., Nguyen, S. C., Proskauer, L., Joyce, E. F., Hasson, D., Jagannathan, M. and Rangan, P. (2024). Genome organization regulates nuclear pore complex formation and promotes differentiation during Drosophila oogenesis. Genes Dev 38, 436–454.

Kotb, N. M., Ulukaya, G., Ramamoorthy, A., Park, L. S., Tang, J., Hasson, D. and Rangan, P. (2025). TORC1-driven translation of Nucleoporin44A promotes chromatin remodeling and germ cell-to-maternal transition in Drosophila. bioRxiv.

Laws, K. M. and Drummond-Barbosa, D. (2015). Genetic Mosaic Analysis of Stem Cell Lineages in the Drosophila Ovary. Methods Mol Biol 1328, 57–72.

Liu, S., Baeg, G. H., Yang, Y., Goh, F. G., Bao, H., Wagner, E. J., Yang, X. and Cai, Y. (2023). The Integrator complex desensitizes cellular response to TGF-β/BMP signaling. Cell Rep 42, 112007.

Liu, W., Chen, H. and King-Jones, K. (2025). Insect nuclear receptors: From orphans to ligands. In Reference Module in Life Sciences: Elsevier.

Losick, V. P., Morris, L. X., Fox, D. T. and Spradling, A. (2011). Drosophila stem cell niches: a decade of discovery suggests a unified view of stem cell regulation. Developmental cell 21, 159–171.

McDonald, S. I., Beachum, A. N., Hinnant, T. D., Blake, A. J., Bynum, T., Hickman, E. P., Barnes, J., Churchill, K. L., Roberts, T. S., Zangwill, D. E., et al. (2019). Novel cis-regulatory regions in ecdysone responsive genes are sufficient to promote gene expression in Drosophila ovarian cells. Gene Expr Patterns 34, 119074.

McKearin, D. M. and Spradling, A. C. (1990). bag-of-marbles: a Drosophila gene required to initiate both male and female gametogenesis. Genes Dev 4, 2242–2251.

Morris, L. X. and Spradling, A. C. (2012). Steroid signaling within Drosophila ovarian epithelial cells sex-specifically modulates early germ cell development and meiotic entry. PLoS One 7, e46109.

Morrow, H. and Mirth, C. K. (2024). Timing Drosophila development through steroid hormone action. Curr Opin Genet Dev 84, 102148.

Nakada, D., Oguro, H., Levi, B. P., Ryan, N., Kitano, A., Saitoh, Y., Takeichi, M., Wendt, G. R. and Morrison, S. J. (2014). Oestrogen increases haematopoietic stem-cell self-renewal in females and during pregnancy. Nature 505, 555–558.

Ng, M., Diaz-Benjumea, F. J. and Cohen, S. M. (1995). Nubbin encodes a POU-domain protein required for proximal-distal patterning in the Drosophila wing. Development 121, 589–599.

Ni, J. Q., Zhou, R., Czech, B., Liu, L. P., Holderbaum, L., Yang-Zhou, D., Shim, H. S., Tao, R., Handler, D., Karpowicz, P., et al. (2011). A genome-scale shRNA resource for transgenic RNAi in Drosophila. Nat Methods 8, 405–407.

Nogueira Alves, A., Oliveira, M. M., Koyama, T., Shingleton, A. and Mirth, C. K. (2022). Ecdysone coordinates plastic growth with robust pattern in the developing wing. Elife 11.

Özel, M. N., Gibbs, C. S., Holguera, I., Soliman, M., Bonneau, R. and Desplan, C. (2022). Coordinated control of neuronal differentiation and wiring by sustained transcription factors. Science 378, eadd1884.

Öztürk-Çolak, A., Marygold, S. J., Antonazzo, G., Attrill, H., Goutte-Gattat, D., Jenkins, V. K., Matthews, B. B., Millburn, G., dos Santos, G., Tabone, C. J., et al. (2024). FlyBase: updates to the Drosophila genes and genomes database. Genetics 227.

Pang, L. Y., DeLuca, S., Zhu, H., Urban, J. M. and Spradling, A. C. (2023). Chromatin and gene expression changes during female Drosophila germline stem cell development illuminate the biology of highly potent stem cells. Elife 12.

Perez-Mockus, G., Cocconi, L., Alexandre, C., Aerne, B., Salbreux, G. and Vincent, J. P. (2023). The Drosophila ecdysone receptor promotes or suppresses proliferation according to ligand level. Developmental cell 58, 2128–2139.e2124.

Peterson, A. J., Murphy, S. J., Mundt, M. G., Shimell, M., Leof, E. B. and O’Connor, M. B. (2022). A juxtamembrane basolateral targeting motif regulates signaling through a TGF-β pathway receptor in Drosophila. PLoS Biol 20, e3001660.

Phipps, D. N., Powell, A. M. and Ables, E. T. (2023). Utilizing the FLP-Out System for Clonal RNAi Analysis in the Adult Drosophila Ovary. Methods Mol Biol 2626, 69–87.

Plavicki, J. S., Squirrell, J. M., Eliceiri, K. W. and Boekhoff-Falk, G. (2016). Expression of the Drosophila homeobox gene, Distal-less, supports an ancestral role in neural development. Dev Dyn 245, 87–95.

Riddiford, L. M. (1993). Hormone receptors and the regulation of insect metamorphosis. Receptor 3, 203–209.

Rørth, P. (1998). Gal4 in the Drosophila female germline. Mech Dev 78, 113–118.

Rust, K., Byrnes, L. E., Yu, K. S., Park, J. S., Sneddon, J. B., Tward, A. D. and Nystul, T. G. (2020). A single-cell atlas and lineage analysis of the adult Drosophila ovary. Nat Commun 11, 5628.

Samuels, T. J., Gui, J., Gebert, D. and Karam Teixeira, F. (2024). Two distinct waves of transcriptome and translatome changes drive Drosophila germline stem cell differentiation. Embo j 43, 1591–1617.

Sanchez, C. G., Teixeira, F. K., Czech, B., Preall, J. B., Zamparini, A. L., Seifert, J. R., Malone, C. D., Hannon, G. J. and Lehmann, R. (2016). Regulation of Ribosome Biogenesis and Protein Synthesis Controls Germline Stem Cell Differentiation. Cell Stem Cell 18, 276–290.

Schauer, S., Callender, J., Henrich, V. C. and Spindler-Barth, M. (2011). The N-terminus of ecdysteroid receptor isoforms and ultraspiracle interacts with different ecdysteroid response elements in a sequence specific manner to modulate transcriptional activity. J Steroid Biochem Mol Biol 124, 84–92.

Schubiger, M., Tomita, S., Sung, C., Robinow, S. and Truman, J. W. (2003). Isoform specific control of gene activity in vivo by the Drosophila ecdysone receptor. Mech Dev 120, 909–918.

Sedkov, Y., Cho, E., Petruk, S., Cherbas, L., Smith, S. T., Jones, R. S., Cherbas, P., Canaani, E., Jaynes, J. B. and Mazo, A. (2003). Methylation at lysine 4 of histone H3 in ecdysone-dependent development of Drosophila. Nature 426, 78–83.

Shi, J., Jin, Z., Yu, Y., Zhang, Y., Yang, F., Huang, H., Cai, T. and Xi, R. (2021). A Progressive Somatic Cell Niche Regulates Germline Cyst Differentiation in the Drosophila Ovary. Current biology : CB 31, 840–852.e845.

Silva, D., Olsen, K. W., Bednarz, M. N., Droste, A., Lenkeit, C. P., Chaharbakhshi, E., Temple-Wood, E. R. and Jemc, J. C. (2016). Regulation of Gonad Morphogenesis in Drosophila melanogaster by BTB Family Transcription Factors. PLoS One 11, e0167283.

Simandi, Z., Cuaranta-Monroy, I. and Nagy, L. (2013). Nuclear receptors as regulators of stem cell and cancer stem cell metabolism. Seminars in cell & developmental biology 24, 716–723.

Slaidina, M., Gupta, S., Banisch, T. U. and Lehmann, R. (2021). A single-cell atlas reveals unanticipated cell type complexity in Drosophila ovaries. Genome Res 31, 1938–1951.

Spradling, A. C., Niu, W., Yin, Q., Pathak, M. and Maurya, B. (2022). Conservation of oocyte development in germline cysts from Drosophila to mouse. Elife 11.

Sun, Z., Nystul, T. G. and Zhong, G. (2023). Single-cell RNA sequencing identifies eggplant as a regulator of germ cell development in Drosophila. EMBO Rep 24, e56475.

Sun, Z., Zeng, Y., Nystul, T. G. and Zhong, G. (2025). Single-cell transcriptomic profiling of <em>bam</em> mutant tumor reveals germline heterogeneity and <em>gcrf1</em> as a modulator in <em>Drosophila</em> germ cells. bioRxiv, 2025.2010.2014.682265.

Takemura, M., Bowden, N., Lu, Y. S., Nakato, E., O’Connor, M. B. and Nakato, H. (2021). Drosophila MOV10 regulates the termination of midgut regeneration. Genetics 218.

Talbot, W. S., Swyryd, E. A. and Hogness, D. S. (1993). Drosophila tissues with different metamorphic responses to ecdysone express different ecdysone receptor isoforms. Cell 73, 1323–1337.

Tan, S. W. S., Yip, G. W., Suda, T. and Baeg, G. H. (2018). Small Maf functions in the maintenance of germline stem cells in the Drosophila testis. Redox Biol 15, 125–134.

Teixeira, F. K., Sanchez, C. G., Hurd, T. R., Seifert, J. R., Czech, B., Preall, J. B., Hannon, G. J. and Lehmann, R. (2015). ATP synthase promotes germ cell differentiation independent of oxidative phosphorylation. Nat Cell Biol 17, 689–696.

Thummel, C. S. (1996). Flies on steroids--Drosophila metamorphosis and the mechanisms of steroid hormone action. Trends in genetics : TIG 12, 306–310.

Uyehara, C. M., Leatham-Jensen, M. and McKay, D. J. (2022). Opportunistic binding of EcR to open chromatin drives tissue-specific developmental responses. Proc Natl Acad Sci U S A 119, e2208935119.

Uyehara, C. M. and McKay, D. J. (2019). Direct and widespread role for the nuclear receptor EcR in mediating the response to ecdysone in Drosophila. Proc Natl Acad Sci U S A 116, 9893–9902.

Uyehara, C. M., Nystrom, S. L., Niederhuber, M. J., Leatham-Jensen, M., Ma, Y., Buttitta, L. A. and McKay, D. J. (2017). Hormone-dependent control of developmental timing through regulation of chromatin accessibility. Genes Dev 31, 862–875.

Van Doren, M., Williamson, A. L. and Lehmann, R. (1998). Regulation of zygotic gene expression in Drosophila primordial germ cells. Current biology : CB 8, 243–246.

Villa-Fombuena, G., Lobo-Pecellín, M., Marín-Menguiano, M., Rojas-Ríos, P. and González-Reyes, A. (2021). Live imaging of the Drosophila ovarian niche shows spectrosome and centrosome dynamics during asymmetric germline stem cell division. Development 148.

Wang, Z. and Lin, H. (2005). The division of Drosophila germline stem cells and their precursors requires a specific cyclin. Current biology : CB 15, 328–333.

Weber, U., Rodriguez, E., Martignetti, J. and Mlodzik, M. (2014). Luna, a Drosophila KLF6/KLF7, is maternally required for synchronized nuclear and centrosome cycles in the preblastoderm embryo. PLoS One 9, e96933.

Weikum, E. R., Liu, X. and Ortlund, E. A. (2018). The nuclear receptor superfamily: A structural perspective. Protein Sci 27, 1876–1892.

Williams, A. E. and Ables, E. T. (2023). Visualizing Fusome Morphology via Tubulin Immunofluorescence in Drosophila Ovarian Germ Cells. Methods Mol Biol 2626, 135–150.

Xie, T. and Spradling, A. C. (1998). decapentaplegic is essential for the maintenance and division of germline stem cells in the Drosophila ovary. Cell 94, 251–260.

Xin, T., Greco, V. and Myung, P. (2016). Hardwiring Stem Cell Communication through Tissue Structure. Cell 164, 1212–1225.

Yamanaka, N., Rewitz, K. F. and O’Connor, M. B. (2013). Ecdysone control of developmental transitions: lessons from Drosophila research. Annu Rev Entomol 58, 497–516.

Zhao, T., Xiao, Y., Huang, B., Ran, M. J., Duan, X., Wang, Y. F., Lu, Y. and Yu, X. Q. (2022). A dual role of lola in Drosophila ovary development: regulating stem cell niche establishment and repressing apoptosis. Cell Death Dis 13, 756.

Zike, A. B., Abel, M. G., Fleck, S. A., DeWitt, E. D. and Weaver, L. N. (2025). Estrogen-related receptor is required in adult Drosophila females for germline stem cell maintenance. Dev Biol 524, 132–143.

