## Supplemental Information for "*Ecdysone Receptor* autonomously controls germ cell differentiation in the *Drosophila* ovary"

**Contents:**

**Supplemental Figures S1 – S9.**

- Figure S1. Driving *UASz-EcR-A* and *UASz-EcR-B1* in all somatic cells induces larval lethality.
- Figure S2. *UASz-EcR-B1* expression in dividing cysts is not sufficient to block germ cell differentiation.
- Figure S3. Expression of cell type-specific genes separates germline and somatic cell clusters.
- Figure S4. Depletion of *tkv* or *dpp* is not sufficient to rescue germ cell differentiation in *UASz-EcR-B1*-expressing cells.
- Figure S5. Pseudotime analysis and single-cell velocity reveal stages of differentiation.
- Figure S6. Cell cycle analysis of undifferentiated germ cells.
- Figure S7. Over-expression of *EcR-B1* in undifferentiated cells delays differentiation prior to down-regulation of *Mov10* and up-regulation of Rbfox1.
- Figure S8. *Eip75B* is necessary and sufficient to repress GSC self-renewal, but is dispensible for germ cell differentiation.
- Figure S9. The co-repressor *Smr* is expressed in undifferentiated germ cells.

**Supplemental Table S1. Key reagents used in this study.**

**Supplemental Table S2 (excel). Differential gene expression between undifferentiated germ cells.**

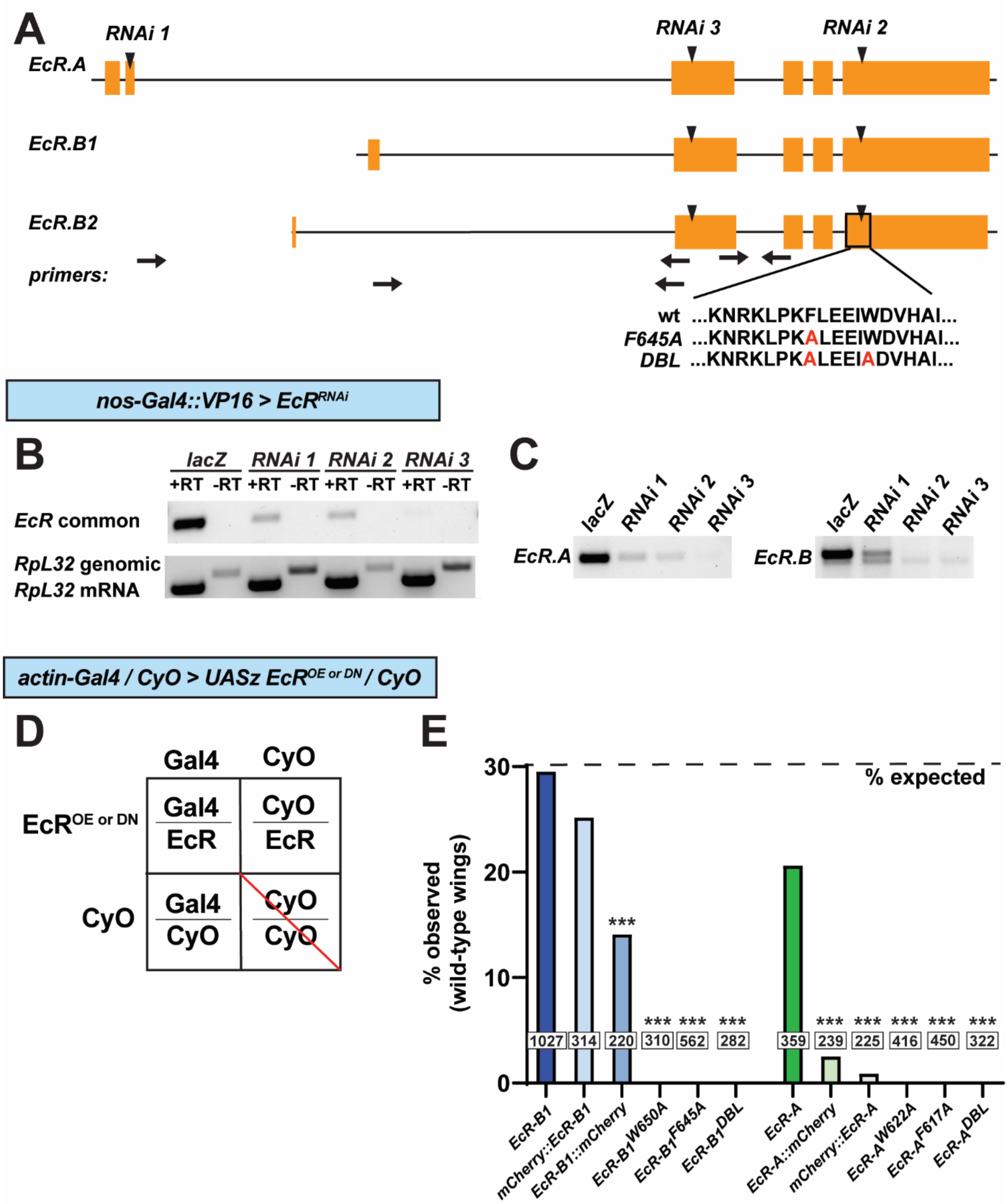

**Figure S1. Development and validation of germline compatible tools to manipulate *EcR*.** (A) The *EcR* gene locus encodes three major mRNA isoforms (*EcR-A, EcR-B1, and EcR-B2*). Arrowheads indicate gene region targeted by RNAi hairpins; arrows indicate gene regions amplified by RT-PCR. (B-C) RT-PCR analysis of RNAi efficiency and isoform specificity in whole ovaries depleted for *EcR* via RNAi hairpins under control of *nos-Gal4::VP16.* (D-E).

*UASz-EcR* transgene efficacy was evaluated by lethality phenotypes under the control of *actin-Gal4*. Mendelian inheritance (D) predicts ~30% expected progeny should carry both *actin-Gal4* and the *UASz-EcR* transgene (dotted line in E). Over-expression of *EcR-B1* or *EcR-A* slightly reduced survival, while expression of the dominant-negative transgenes resulted in nearly complete lethality, consistent with previous studies.

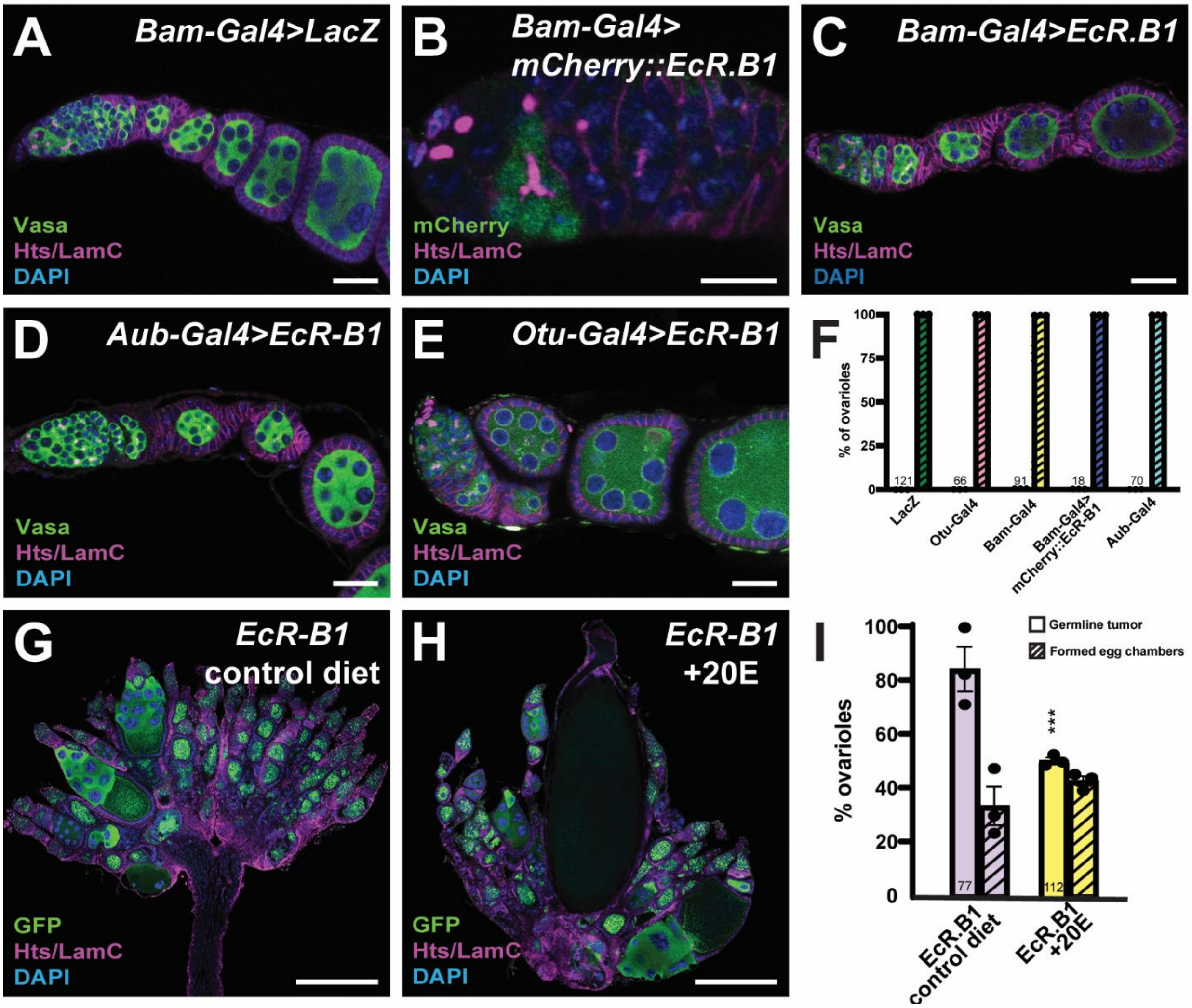

**Figure S2.** ***UASz-EcR-B1*** **expression in dividing cysts is not sufficient to block germ cell differentiation. (A-F)** Representative images (A-E) and quantification (F) of *3x-Bam-Gal4* driving *UASp-lacZ* alone (A, control) or with *UASz-EcR-B1* (B), or *UASz-mCherry::EcR-B1* (C), or from Aub-Gal4 (D) or Otu-Gal4::VP16 (E) driving *UASz-EcR-B1* germaria immunostained for Vasa (green, germ cells, A,C-E) or mCherry (green, transgene expression, B), Hts (magenta; fusomes and follicle cells), LamC (magenta; cap cells and stalk cells), and DAPI (nuclei). In contrast to *nos-Gal4::VP16* (see Figure 2), over-expression of EcR-B1 in cystoblasts or dividing cysts did not block germ cell differentiation. **(G-I)** Representative images (G-H) and quantification (I) of *nos-Gal4::VP16>UASz-EcR-B1* ovaries from females maintained on a control wet yeast diet (G) or wet diet plus excess ecdysone (20E; H). In the presence of excess dietary ecdysone, germaria contained more dividing cysts (4-cell, 8-cell, etc.), ovaries contained more normal egg chambers, and some large eggs ready for fertilization could be detected. Scale bars = 20 μm (except panel B, =10 μm). Bars in (F & I) represent the percentage of ovarioles displaying a block to germ cell differentiation; *n* > 50 ovarioles from at least 15 flies per genotype, ~5 days after eclosion. ***$p$<0.001, compared to control; Fisher's exact t-test.

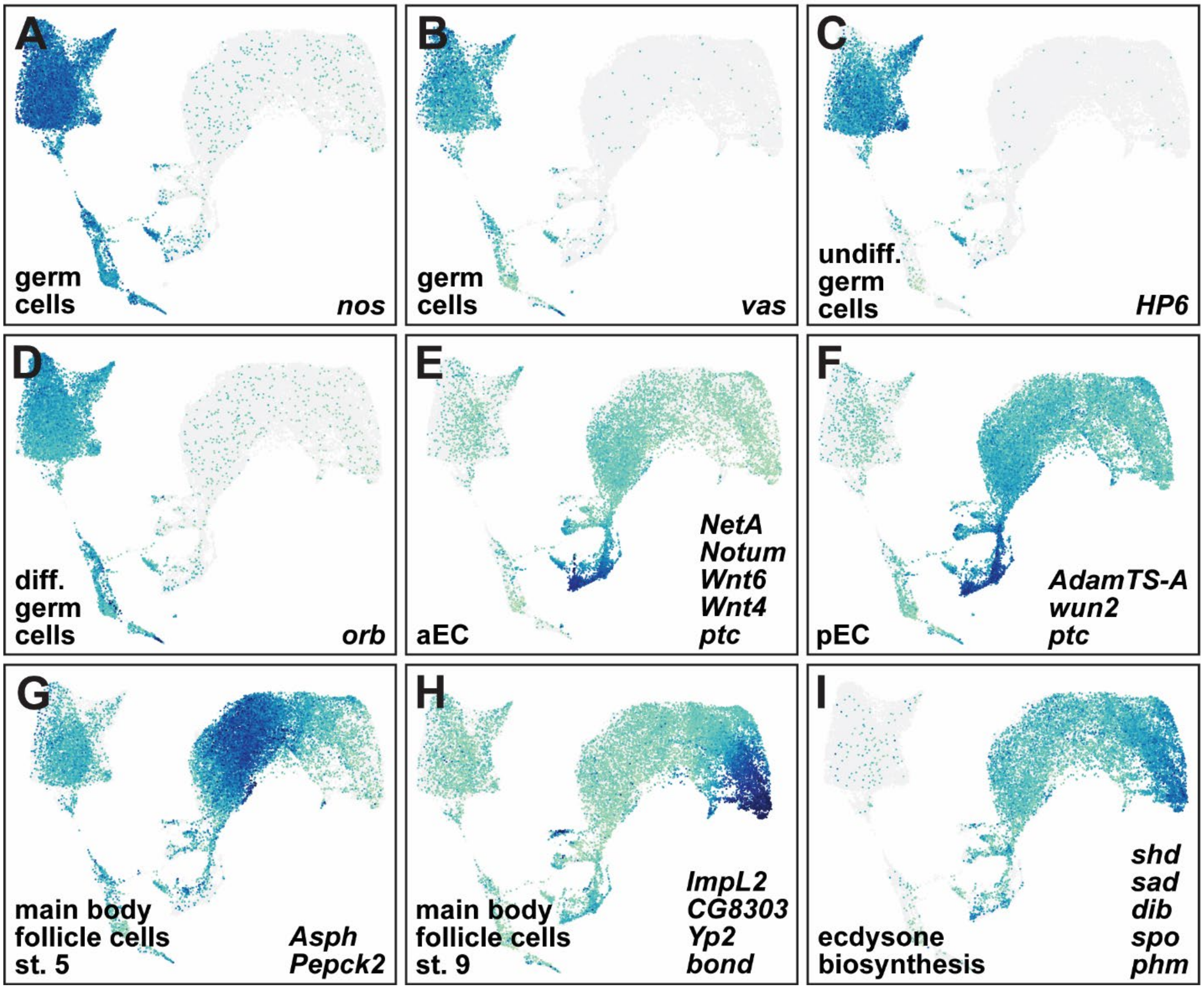

**Figure S3. Expression of cell type-specific genes separates germline and somatic cell clusters.** Representative expression level UMAPs of all cells merged from wildtype, *nos-Gal4::VP16>UASz-EcR-B1*, *nos-Gal4::VP16>UASp-tkv$^{ACT}$*, and *bam$^{\Delta 86}$* ovaries, visualizing: the germ cell markers *nos* (A), *vasa* (B), *HP6* (C), and *orb* (D); the anterior escort cell markers *NetA*, *Notum*, *Wnt4*, *Wnt6*, and *ptc* (E); the posterior escort cell markers *AdamTS-A*, *wun2*, and *ptc* (F); the main body follicle cell markers *Asph* and *Pepck2* (G; stage 5 egg chambers) or *ImpL2*, *CG8303*, *Yp2*, and *bond* (H; stage 9 egg chambers); and the ecdysone biosynthesis genes *shd*, *sad*, *dib*, *spo* and *phm* (I).

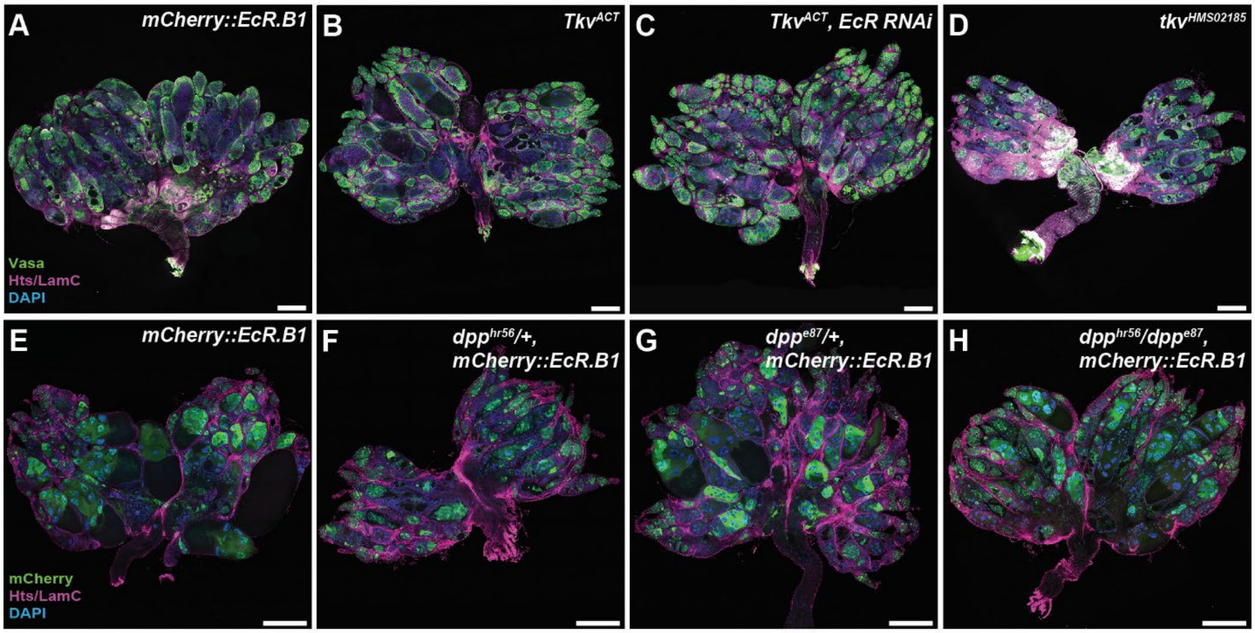

**Figure S4. Depletion of *tkv* or *dpp* is not sufficient to rescue germ cell differentiation in *UASz-EcR.B1*-expressing cells.** (A-D) Knock-down of *tkv* in *EcR* over-expressing germ cells produces equivalent phenotypes as over-expression of *EcR* alone (A), over-expression of t*kv*$^{ACT}$ alone (B), or *tkv* RNAi alone (D). Similarly, reduction of one copy (F-G) or two copies (H) of *dpp* in *EcR* over-expressing germ cells produces equivalent phenotypes as over-expression of *EcR* alone (E). Ovaries were immunostained for Vasa (green, germ cells, A-D) or mCherry (green, transgene expression, E-H), Hts (magenta; fusomes and follicle cells), LamC (magenta; cap cells and stalk cells), and DAPI (nuclei). Scale bars represent 50µm.

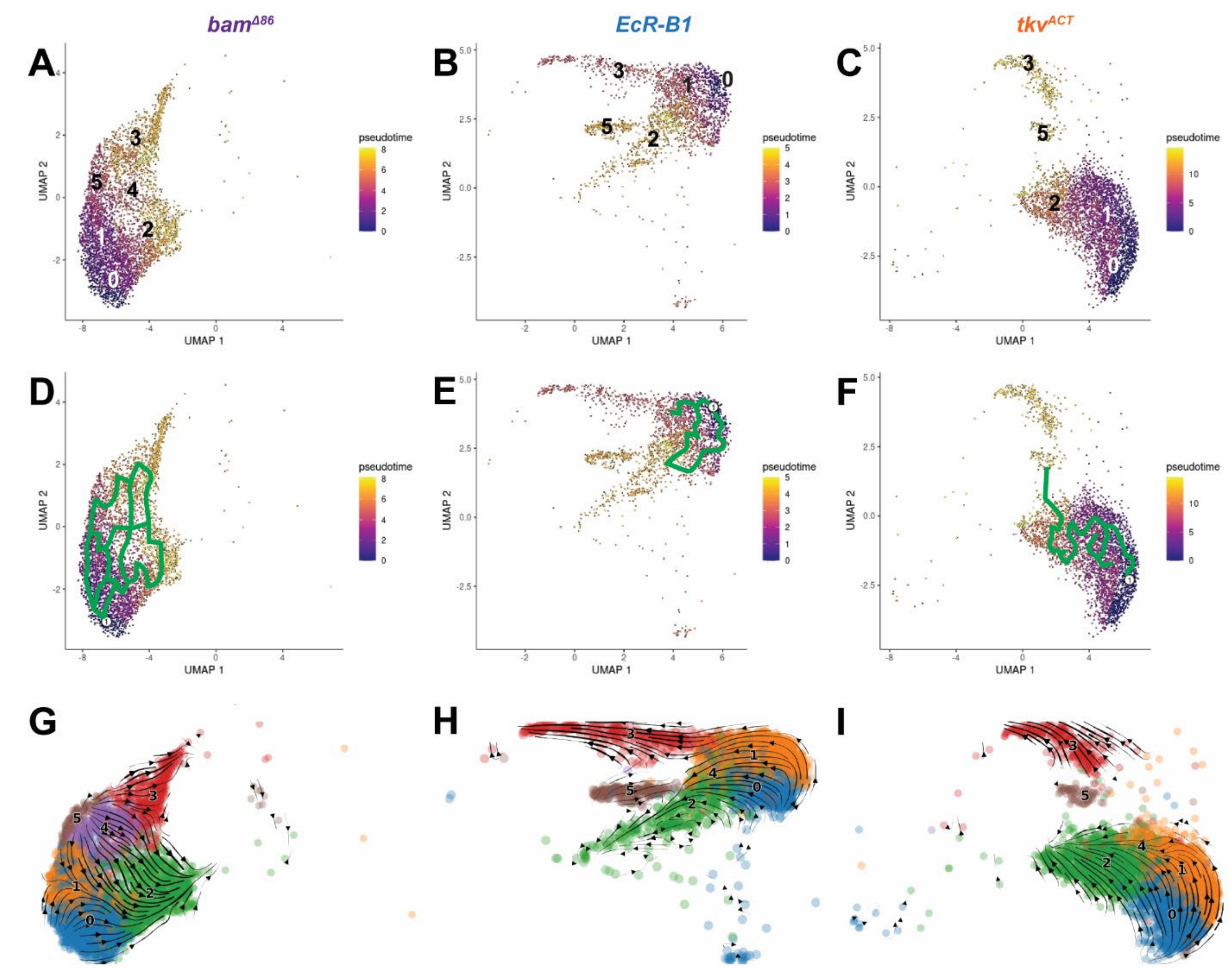

**Figure S5. Pseudotime analysis and single-cell velocity reveal stages of differentiation.** (A-F) Pseudotime analysis of undifferentiated germ cells from (A) $bam^{\Delta 86}$, (B) *EcR-B1*, and (C) $tkv^{ACT}$ ovaries. The lighter the color of the cells, the more differentiated they are. In panels D-F, the estimated trajectory of differentiation is overlayed in green. (G-I) Single cell velocity analysis of (D) $bam^{\Delta 86}$, (E) *EcR-B1*, and (F) $tkv^{ACT}$. Undifferentiated germ cell subtype 0 is in blue, 1 in orange, 2 in green, 3 in red, 4 in purple, and 5 in brown.

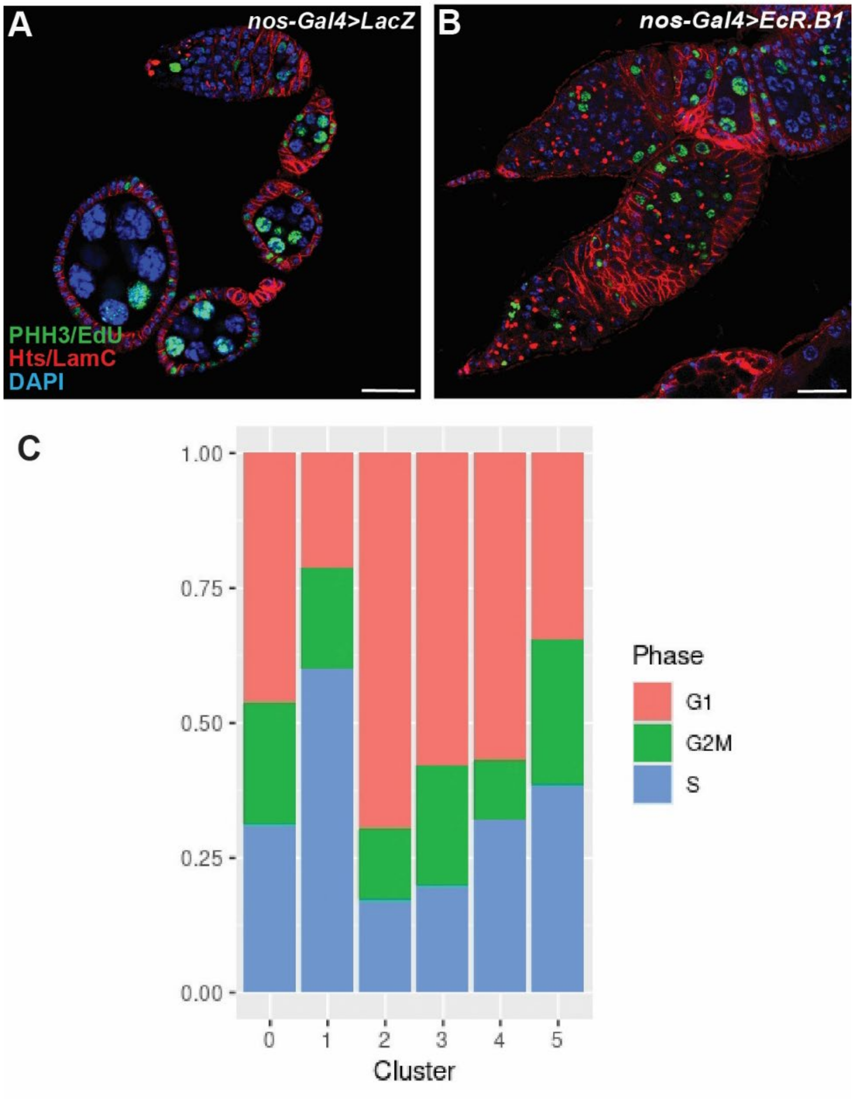

**Figure S6. *EcR* over-expressing germ cells are proliferative.** (A-B) Control (A) and EcR over-expression (B) ovaries immunostained for the active cell cycle markers phosphorylated Histone H3 (PHH3) and EdU (green), Hts (magenta; fusomes and follicle cells), LamC (magenta; cap cells and stalk cells), and DAPI (nuclei). Scale bar = 20 μM. (C) Cell cycle phase estimation of single undifferentiated germ cells from wildtype, *nos-Gal4::VP16>UASz-EcR-B1*, *nos-Gal4::VP16>UASp-tkv*$^{ACT}$, and *bam*$^{\Delta 86}$ ovaries, based on CellCycleScoring (Zhu et al., 2022) of cell cycle transcripts in single cell mRNA datasets. Numbers at the bottom of each column represent undifferentiated germ cell subtype (0-5).

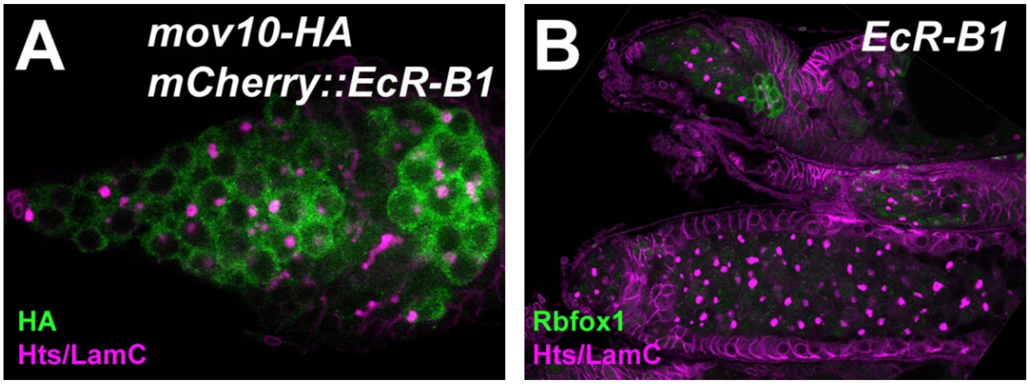

**Figure S7. Over-expression of EcR-B1 in undifferentiated cells delays differentiation prior to down-regulation of Mov10 and up-regulation of Rbfox1.** *nos-Gal4::VP16>UASz-EcR-B1, mov10*$^{KI\text{-}HA}$ ovaries immunostained for HA (A; *Mov10* reporter; green) or Rbfox1 (B; green), Hts (magenta; fusomes and follicle cells), and LamC (magenta; cap cells and stalk cells).

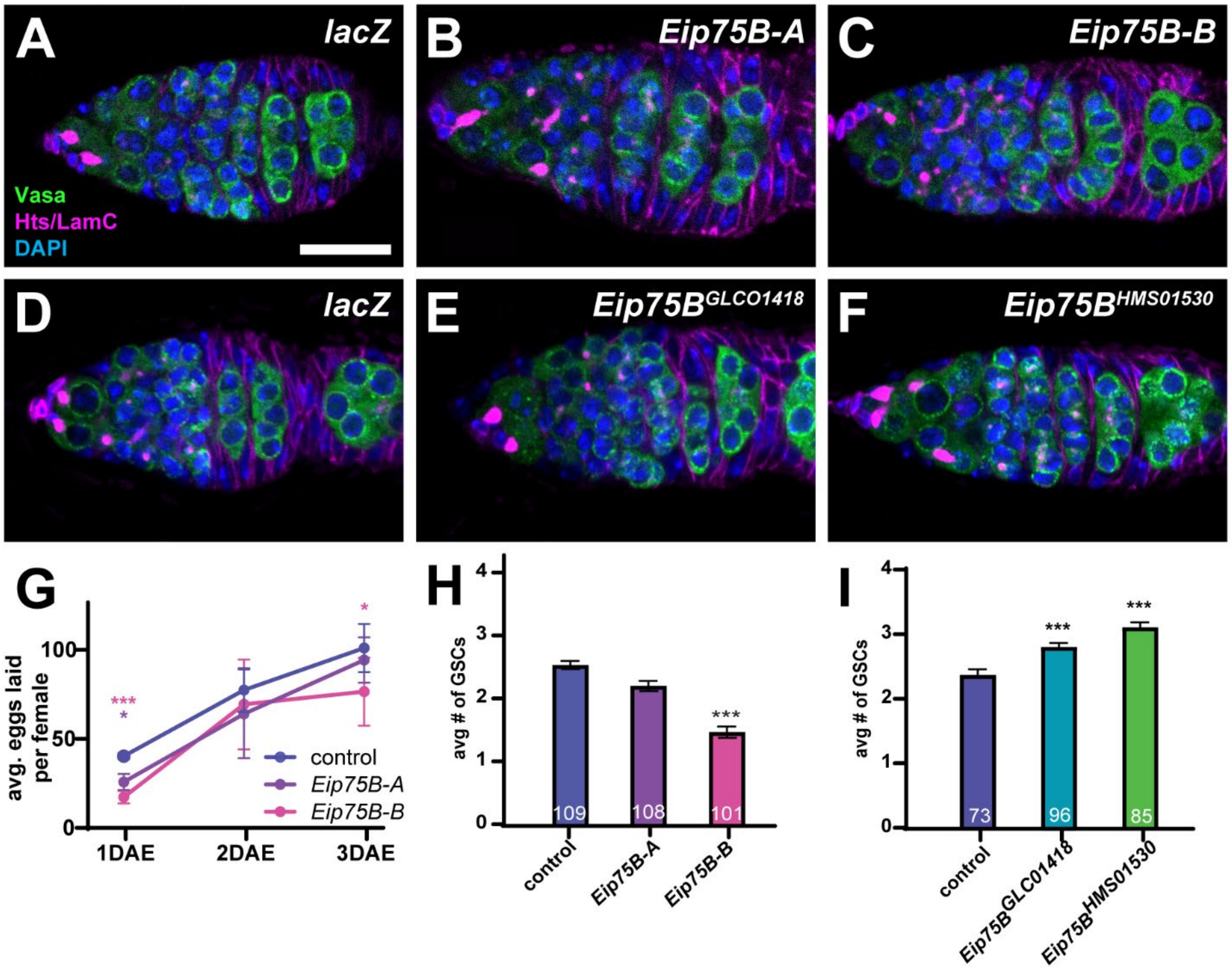

**Figure S8. *Eip75B* is necessary and sufficient to repress GSC self-renewal, but is dispensible for germ cell differentiation.** (A-F) Representative images (A-F) and quantification (G-I) of control (A,D), *nos-Gal4::VP16>UASz-Eip75B* over-expression (B-C), or depletion of *Eip75B* via germline-enhanced RNAi ($Eip75B^{GLC01418}$ and $Eip75B^{HMS01530}$) germaria immunostained for Vasa (germ cells; green), Hts (magenta; fusomes and follicle cells), LamC (magenta; cap cells and stalk cells), and DAPI (nuclei). Scale bars = 10 µm. (G) Average number of eggs laid per female over the first three days after eclosion (DAE). (H-I) Average number of GSCs per germarium at 5 days after eclosion. Numbers in bars represent number of germaria scored. **p*<0.01, ****p*<0.001, compared to control; Student's t-test.

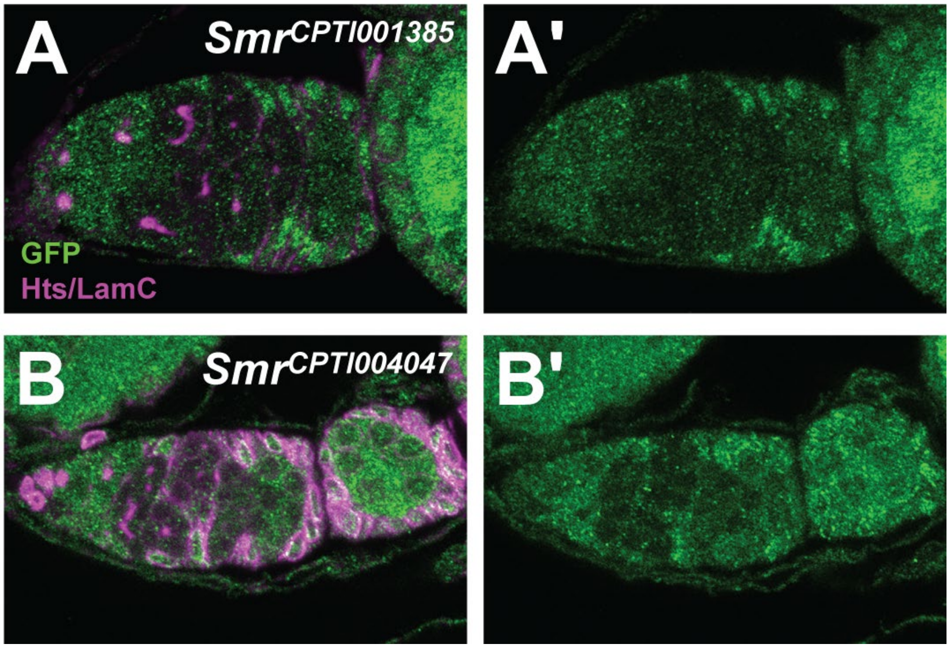

**Figure S9. The co-repressor *Smr* is expressed in undifferentiated germ cells.** Flies carrying GFP-tagged protein trap *Smr* reporters (A-A', $Smr^{CPTI001385}$; B-B', $Smr^{CPTI004047}$) immunostained for GFP (reporter expression; green), Hts (magenta; fusomes and follicle cells), and LamC (magenta; cap cells and stalk cells).

### Table S1. Reagents used in this study.

| ***Drosophila* Stocks** | | |
|---|---|---|
| **Stock Name** | **Source** | **Identifier** |
| *UASp-lacZ* | D. Drummond-Barbosa | Bloomington *Drosophila* Stock Center (BDSC) #98113 |
| $w^{1118}$; P{GAL4::VP16-nanos.UTR}$CG6325^{MVD1}$ (nos-Gal4::VP16) | D. Drummond-Barbosa (Rørth, 1998; Van Doren et al., 1998) | BDSC #4937 or #7253 |
| $w^{*}$; P{Act5C-GAL4}25FO1/CyO; M{3xP3-GFP.R}ZH-86Fb | BDSC | BDSC #78574 |
| $y^{1}$ $w^{*}$; TI{CRIMIC.TG4.1}$aub^{CR02423-TG4.1}$/SM6a (Aub-Gal4) | BDSC | BDSC #92663 |
| P{otu-GAL4::VP16.R}1, $w^{1118}$ | BDSC | BDSC #58424 |
| *bam>Gal4^A,Bam>Gal4^C/CyO;Bam>Gal4/TM6 (3xbam-Gal4)* | J. Mathieu & J.-R. Huynh (Clemot et al 2018) | |
| UASz-EcR-$B1^{\Delta C655}$ (attP40)/CyO | this manuscript | |
| UASz-mCherry::EcR-$B1^{\Delta C655}$ (attP40)/CyO | this manuscript | |
| UASz-mCherry::EcR-$B1^{\Delta C655}$ (attP2)/TM6 | this manuscript | |
| UAS-$EcR^{RNAi1\ (Valium22-attP2)}$/TM3 | this manuscript | |
| UAS-$EcR^{RNAi2\ (Valium22-attP2)}$/TM3 | this manuscript | |
| UAS-$EcR^{RNAi3\ (Valium22-attP2)}$/TM3 | this manuscript | |
| *UASz-HA::Eip75B-A (attP2) / TM3* | this manuscript | |
| *UASz-HA::Eip75B-B (attP2) / TM3* | this manuscript | |
| UASp-$tkv^{Q199D}$/TM6 | D. Drummond-Barbosa (Bolivar et al., 2006) | |
| $ry^{506}$ $e^{1}$ $bam^{\Delta 86}$/TM3, $ry^{RK}$ $Sb^{1}$ $Ser^{1}$ | BDSC | BDSC #5427 |
| $y^{1}$ $v^{1}$; P{TRiP.HMJ22155}attP40 (bam RNAi) | BDSC | BDSC #58178 |
| $y^{1}$ $v^{1}$; P{TRiP.HMS02185}attP40 (tkv RNAi) | BDSC | BDSC #40937 |
| $y^{1}$ $sc^{*}$ $v^{1}$ $sev^{21}$; P{TRiP.HMS04501}attP40 (tkv RNAi) | BDSC | BDSC #57303 |
| $dpp^{hr56}$ $cn^{1}$ $bw^{1}$/CyO | BDSC | BDSC #36528 |
| $dpp^{e87}$ $cn^{1}$ $bw^{1}$/CyO | BDSC | BDSC #2073 |
| *PBac{fTRG00003.sfGFP-TVPTBF}VK00033 (Bam::GFP)* | Vienna Drosophila Resource Center | v318001 |
| Dad-lacZ (likely P{lacZ}$Dad^{P1883}$) | D. Drummond-Barbosa (Kai and Spradling, 2003; Song et al., 2004) | |
| $y^{1}$ $w^{*}$; Mi{PT-GFSTF.1}Eip63EMI00413-GFSTF.1/TM6C, $Sb^{1}$ $Tb^{1}$ | BDSC | BDSC #59763 |
| *w; dMov10[KI, smGFP-HA]/SM; Pr.e/TM3 Ser* | H. Nakato (Takemura et al., 2021) | |
| $y^{1}$ $w^{*}$; Mi{MIC}$Eip93F^{MI05200}$/TM3, $Sb^{1}$ $Ser^{1}$ | BDSC | BDSC #43675 |

| | | |
|---|---|---|
| $y^1$ $w^*$; P{E(bx)-GFP.FPTB}attP40 | BDSC | BDSC #68180 |
| $y^1$ $w^*$; PBac{crol-GFP.FPTB}VK00031 | BDSC | BDSC #93087 |
| $w^{1118}$; PBac{HmgD-GFP.FPTB}VK00033 | BDSC | BDSC #55827 |
| $w^{1118}$; PBac{Hr78-GFP.FLAG}VK00037 | BDSC | BDSC #38653 |
| $w^{1118}$; PBac{Hnf4-GFP.FLAG}VK00033 | BDSC | BDSC #38649 |
| $y^1$ $w^*$; PBac{Dp-GFP.FPTB}VK00033 | BDSC | BDSC #67388 |
| *w*; P{UASp-hts.mCherry}attP2* | BDSC | BDSC #66171 |
| $w^{1118}$; PBac{lola.DE-GFP.FLAG}VK00033/TM3, $Sb^1$ | BDSC | BDSC #43948 |
| $w^{1118}$; PBac{lola.I-GFP.FLAG}VK00033 | BDSC | BDSC #38662 |
| $y^1$ $w^{67c23}$; $ftz\text{-}f1^{ex7}$ P{FRT($w^{hs}$)}2A/TM6B, P{Ubi-GFP.S65T}PAD2, $Tb^+$ | BDSC | BDSC #64338 |
| $w^*$; P{FRT($w^{hs}$)}2A (FRT79D) | BDSC | BDSC #1997 |
| *hsFLP; nGFP FRT79D/TM3* | M. Buszczak | |
| *UASz-3xHA::Eip75B-A (attP2)/TM3 Sb* | this manuscript | |
| *UASz-3xHA::Eip75B-B (attP2)/TM3 Sb* | this manuscript | |
| $w^{1118}$ PBac{806.LOX-SVS-2}$Smr^{CPTI004047}$ | Kyoto *Drosophila* Stock Center | 115465 |
| $w^{1118}$ PBac{681.P.FSVS-1}$Smr^{CPTI001385}$ | Kyoto *Drosophila* Stock Center | 115513 |
| **Antibodies and Stains** | | |
| **Antibody** | **Source** | **Identifier** |
| Mouse anti-Hts (1:10) | Developmental Studies Hybridoma Bank (DSHB) | 1B1; AB_528070 |
| Mouse anti-LamC (1:100) | DSHB | LC28.26; AB_528339 |
| Chicken anti-GFP (1:2000) | Abcam | AB_13970 |
| Rabbit anti-Vasa (1:1000) | P. Lasko (Lasko & Ashburner et a., 1990) | |
| Rabbit anti-Vasa (1:1000) | Boster Biological | DZ41154 |
| Chicken anti-Vasa (1:5000) | P. Rangan (Upadhyay et al., 2016) | |
| Chicken anti-β gal | Abcam | AB_9361 |
| Rabbit anti-DsRed (1:500) | Takara | 632496 |
| Rabbit anti-Tkv2 (Tkv-A) (1:250) | M. O'Connor (Peterson et al., 2022) | |
| Guinea pig anti-Rbfox1 (1:5000) | M. Buszczak (Tastan et al 2010) | |
| Rabbit anti-pMad (1:100) | Abcam | AB_52903 |
| Mouse anti-Orb (1:500) | DSHB | 4H8; AB_528418 |
| Mouse anti-Orb (1:500) | DSHB | 6H4; AB_528419 |
| Rabbit anti-Blanks (1:1000) | Erik Sontheimer (Gerbasi et al. 2011) | |

| | | |
|---|---|---|
| Rat anti-Chinmo (1:1000) | Nick Sokol and Lesley Weaver (Wu et al 2012) | |
| Mouse anti-Lola (1:20) | DSHB | 7F1-1D5 RRID: AB_2721954 |
| Rabbit anti-pHistone H3 (pHH3) (1:200) | Millipore | 06-570 |
| Click-iT EdU Cell Proliferation Kit for Imaging, Alexa Fluor 647 dye | Life Technologies | C10340 |
| Goat anti-mouse AlexaFluor 488, 568, 633 (1:200) | Life Technologies | |
| Goat anti-chicken AlexaFluor 488 (1:200) | Life Technologies | |
| Goat anti-rabbit AlexaFluor 488, 568 (1:200) | Life Technologies | |
| Goat anti-rat Alexa Fluor 488 (1:200) | Life Technologies | |
| 4'-6-diamidino-2-phenylindole (DAPI) (0.5 μg/mL) | Sigma | |
| **Primers** | | |
| **Primer Name** | **Sequence (5'-3')** | |
| UASp-Tkv$^{ACT}$ forward | GTCATCAAGCTTAGGCCTCCAA | |
| UASp-Tkv$^{ACT}$ reverse | TCCCGGTCGTCTCATCGTAA | |
| pVALIUM 22 forward | GGT GAT AGA GCC TGA ACC AG | |
| pVALIUM 22 reverse | TAA TCG TGT GTG ATG CCT ACC | |
| UASz-EcR.B1 C Term forward | GCTGTACAAGATGAAGCGGCGCTGGTCG | |
| UASz-EcR.B1 C Term reverse | AGTGGTACCCTCGAGGGATCCTAGATGGCATGAACGTCCCAG | |
| EcR common forward (exon 3-4) | CCTCCGGCTACCACTACAAC | |
| EcR common reverse | TATTGCGCGCTTGACACTTG | |
| EcR-A forward (exon 2-3) | CGCCGGAAGCTATGATCCTT | |
| EcR-A reverse | TCCTTCTCCTTCTGGGCCTT | |
| EcR-B forward (exon 1-3) | ATGGCCTAAGTCAGCAGCAG | |
| EcR-B forward | CTTGCTCTTCTTCGCATCGC | |
